# Amplifying the redistribution of somato-dendritic inhibition by the interplay of three interneuron types

**DOI:** 10.1101/410340

**Authors:** Loreen Hertäg, Henning Sprekeler

## Abstract

GABAergic interneurons play an important role in shaping the activity of excitatory pyramidal cells (PCs). How the various inhibitory cell types contribute to neuronal information processing, however, is not resolved. Here, we propose a functional role for a widespread network motif consisting of parvalbumin-(PV), somatostatin-(SOM) and vasoactive intestinal peptide (VIP)-expressing interneurons. Following the idea that PV and SOM interneurons control the distribution of somatic and dendritic inhibition onto PCs, we suggest that mutual inhibition between VIP and SOM cells translates weak inputs to VIP interneurons into large changes of somato-dendritic inhibition of PCs. Using a computational model, we show that the neuronal and synaptic properties of the circuit support this hypothesis. Moreover, we show that the SOM-VIP motif allows transient inputs to persistently switch the circuit between two processing modes, in which top-down inputs onto apical dendrites of PCs are either integrated or canceled.

## 1. Introduction

GABAergic interneurons are essential for maintaining normal brain activity (Isaacson and Scanziani, 2011; Marín, 2012; Tremblay et al., 2016), although they are outnumbered by excitatory cells throughout the brain (Meyer et al., 2011). They present a large number of distinct types that differ in their anatomical, physiological and biophysical properties (Markram et al., 2004; Gentet, 2012; Tremblay et al., 2016; Naka and Adesnik, 2016; Wamsley and Fishell, 2017). This has led to the hypothesis that individual types are optimized to perform specific computations in neuronal microcircuits (Silberberg and Markram, 2007; Adesnik et al., 2012; Kvitsiani et al., 2013; Hangya et al., 2014; Kepecs and Fishell, 2014; Tremblay et al., 2016). The functional roles of these interneuron classes and how they are supported by their individual characteristics, however, are still largely unknown.

One conspicuous difference between interneuron types is the location of their synapses onto their postsynaptic targets: Parvalbumin-expressing (PV) interneurons preferably inhibit the perisomatic regions and the basal dendrites of excitatory pyramidal cells (PCs), as well as other PV neurons (Rudy et al., 2011; Avermann et al., 2012; Pfeffer et al., 2013; Hu et al., 2014; Jiang et al., 2015; Tremblay et al., 2016). In contrast, somatostatin-expressing (SOM) neurons mainly target the apical dendrites of PCs, and strongly inhibit other interneuron types (Pfeffer et al., 2013; Jiang et al., 2015; Yavorska and Wehr, 2016; Urban-Ciecko and Barth, 2016). A third group expressing vasoactive intestinal peptide (VIP) mainly connects to the dendrite-targeting SOM neurons. In addition to these distinct connectivity motifs, different interneuron types also differ in their intrinsic and synaptic properties. For instance, PV neurons hardly exhibit spike-frequency adaptation (Rudy et al., 2011; Hu et al., 2014; Tremblay et al., 2016), a neuronal characteristic that has been observed both for SOM (Urban-Ciecko and Barth, 2016; Tremblay et al., 2016) and VIP cells (Tremblay et al., 2016).

As a consequence of the interneuron-specific, spatially distinct distribution of synapses onto PCs and the direct connection from SOM to PV neurons, it has been hypothesized that the SOM-PV motif plays a key role in the redistribution of somatic and dendritic inhibition (Pouille and Scanziani, 2004; Pfeffer et al., 2013). Inhibiting specific compartments of PCs may have wide-ranging functional and computational consequences, because their somata and dendrites are the target of two distinct information streams. Top-down input originating from higher cortical areas and non-specific thalamocortical pathways (Felleman and Van, 1991; Cauller et al., 1998; Diamond, 1995) selectively aims at apical dendrites (Larkum, 2013). At the same time, feedforward bottom-up input from lower cortical areas and the core thalamic nuclei arrives at perisomatic regions and basal dendrites (Larkum, 2013). While top-down feedback is associated with internal predictions, bottom-up connections are thought to carry information from the external world (Larkum, 2013). Hence, control of the different input streams – and consequently, information processing modes – is of fundamental importance.

Here, we hypothesize that a different subnetwork consisting of SOM and VIP neurons is optimized to efficiently control the PV/SOM-mediated redistribution of somatic and dendritic inhibition. In order to support our hypothesis, we perform mathematical analyses and extensive simulations of a microcircuit consisting of these three interneuron types and excitatory PCs. We show that mutual inhibition between SOM and VIP cells leads to an amplification of weak signals targeting VIP neurons, which in the extreme case of strong mutual inhibition turns the SOM-VIP motif into a switched-mode amplifier. Furthermore, we reveal how frequently reported connectivity, neuronal and synaptic properties underpin the amplification abilities of the microcircuit, such as the lack of recurrent connections among both SOM and VIP cells, their prominent spike-frequency adaptation and short-term facilitation. Moreover, we show that the circuit can display slow oscillations ranging from Delta to Alpha bands as a consequence of spike-frequency adaptation and strong mutual inhibition in SOM and VIP neurons.

Functionally, strong mutual inhibition between SOM and VIP neurons enables a switch between two distinct processing modes in which top-down inputs arriving at the apical dendrites of PCs are either integrated or obliterated via VIP cell modulation. The transition between these operating modes can be triggered by either weak and persistent input or strong and transient pulses.

## 2. Results

We study a rate-based network model consisting of excitatory PC and inhibitory PV, SOM and VIP cells (see Figure 1 A). The ratio of excitatory and inhibitory neurons and the strength and probability of their connections are constrained by experimental findings (Bartley et al., 2008; Fino and Yuste, 2011; Packer and Yuste, 2011; Avermann et al., 2012; Pfeffer et al., 2013; Lee et al., 2013; Pi et al., 2013; Jiang et al., 2015; Jouhanneau et al., 2015; Pala and Petersen, 2015; Urban-Ciecko et al., 2015, see Tables 1-3 in Models and Methods). While GABAergic neurons are described by point neuron models (Wilson and Cowan, 1972), PCs are modeled as two compartments, to capture both somatic activity and active processes in their apical dendrites (Murayama et al., 2009, see Models and Methods).

**Figure 1.**
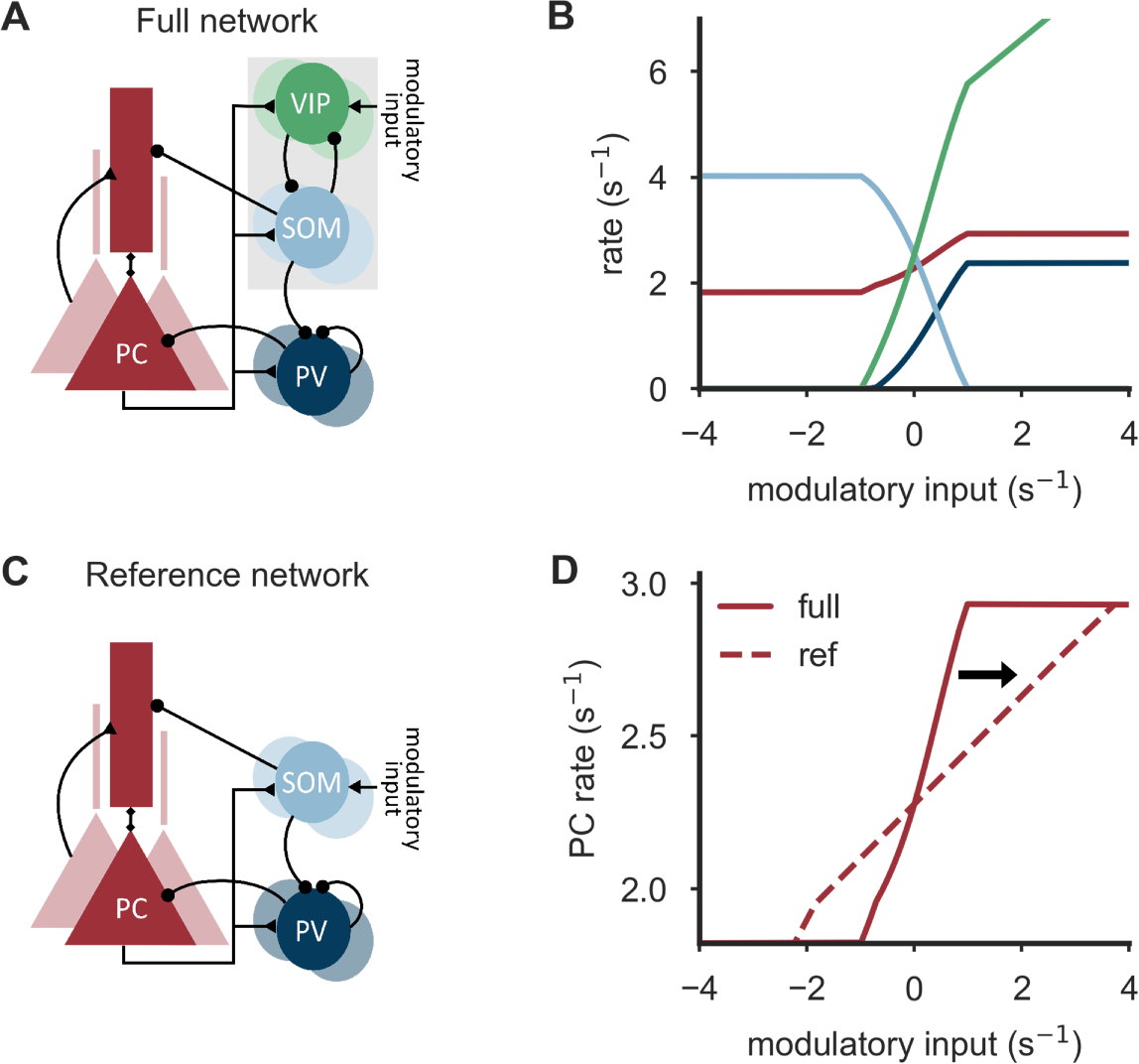
Amplifying the redistribution of inhibition along PC neurons by the SOM-VIP motif. **(A)** Connectivity of the circuit model, inspired by experimentally observed connectivity of excitatory pyramidal cells (PCs) and inhibitory PV, SOM and VIP neurons (see main text). VIP neurons receive an additional, modulatory input. **(B)** Population rates of all neuron types as a function of the modulatory input onto VIP cells. The PC rate follows a sigmoid function, the slope of which characterizes the gain of the redistribution of somato-dendritic inhibition upon a change in the modulatory input. **(C)** Reference network without VIP neurons, in which modulatory input targets SOM neurons instead. **(D)** The firing rate curve of the PCs in the full network (A) exhibit a larger slope than in the reference network without VIP neurons (C). Parameters (A-D): Mutual inhibition strength 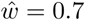 = 0.7, adaptation strength *b* = 0.2, initial synaptic efficacy *U*_s_ = 0.4.

All neurons receive background input to ensure similar firing rates as observed *in vivo* (see e.g. Gentet, 2012; Pala and Petersen, 2015; Urban-Ciecko and Barth, 2016). PC input is divided into two separate information streams: top-down feedback arriving at the apical dendrite and bottom-up input targeting the perisomatic region.

VIP cells receive an additional, modulatory input that regulates the distribution of somatic and dendritic inhibition onto PCs as follows (see Figure 1 B): When the modulatory VIP input is sufficiently small or even inhibitory, VIP neurons remain inactive. As this relieves the SOM neurons from VIP inhibition, they can in turn inhibit the apical dendrites of the PCs and thereby suppress potential top-down inputs. At the same time, the amount of somatic inhibition in PCs is reduced, because SOM cells inhibit PV neurons. Once VIP cells are fully deactivated, further reducing the modulatory input has no effect on the PCs, as the modulatory input acts through VIP neurons only (Figure 1 A). The opposite scenario is a strong and excitatory modulatory input that renders VIP cells sufficiently active to silence SOM neurons. Silencing SOM cells removes dendritic inhibition, so that PCs are receptive to both bottom-up input and top-down feedback. In turn, the perisomatic compartments of PCs experience more inhibition, because PV neurons are released from SOM neuron inhibition. Once the VIP cells are sufficiently active to silence SOM neurons, further increasing the modulatory input has no effect on the PCs, because VIP cells act through SOM neurons only (Figure 1 A). VIP neurons then effectively decouple from the microcircuit. In between these two extremes of inactive VIP or SOM neurons, respectively, the ratio of somatic and dendritic inhibition can be controlled by adjusting the modulatory signal. This is reflected by the relationship between modulatory VIP input and PC activity, which resembles a sigmoid function (Figure 1 B), the slope of which characterizes the gain of the redistribution of somatic and dendritic inhibition upon a change in the modulatory input.

In principle, a similar somato-dendritic redistribution of inhibition could also be achieved by modulatory input directly to SOM neurons. To understand the role of VIP neurons in the circuit, we considered a ‘reference network’ without VIP neurons (Figure 1 C), in which the modulatory input targets SOM cells instead (with inverted sign for comparability). We found that in this reference network, the slope of the corresponding sigmoid function decreases (Figure 1 D) for a large parameter range, indicating that inputs onto VIP neurons in the SOM-VIP motif are more effective modulators than inputs onto SOM neurons. This observation led us to the hypothesis that the SOM-VIP motif serves to translate weak signals onto VIP neurons into large changes of the somatodendritic distribution of inhibition. We therefore wondered whether the connectivity and the neuronal and synaptic properties of the circuit are optimized to support this function, and which computational purpose the circuit could fulfill. To address these questions, computational modeling is well suited, because it allows to study the effect of arbitrary manipulations and variations of the circuit.

### Mutual inhibition between SOM and VIP neurons creates an amplifier

To gain a deeper understanding of the circuit mechanisms and the interplay of the interneuron types, we next studied a simplified microcircuit consisting only of the three interneuron classes expressing PV, SOM and VIP (Figure 2 A, left). The advantage of this simpler model is that it bypasses the nonlinearities of dendritic integration in PCs, thereby allowing an in-depth mathematical analysis of the parameter dependences of the network. All results are later verified in the full circuit. In the simplified network, the local PC input onto the GABAergic interneurons is replaced by additional excitatory inputs to maintain realistic firing rates. Compartment-specific inhibition onto PCs is represented by the population firing rate of the respective interneuron type: the rate of SOM neurons reflects the strength of dendritic inhibition and the rate of PV neurons the strength of somatic inhibition. Similar to the PC rate in the full microcircuit, the difference of the PV and SOM neuron rates shows a sigmoidal dependence on the modulatory VIP cell input (Figure 2 B), whose slope quantifies the system’s sensitivity to changes in the modulatory input.

**Figure 2.**
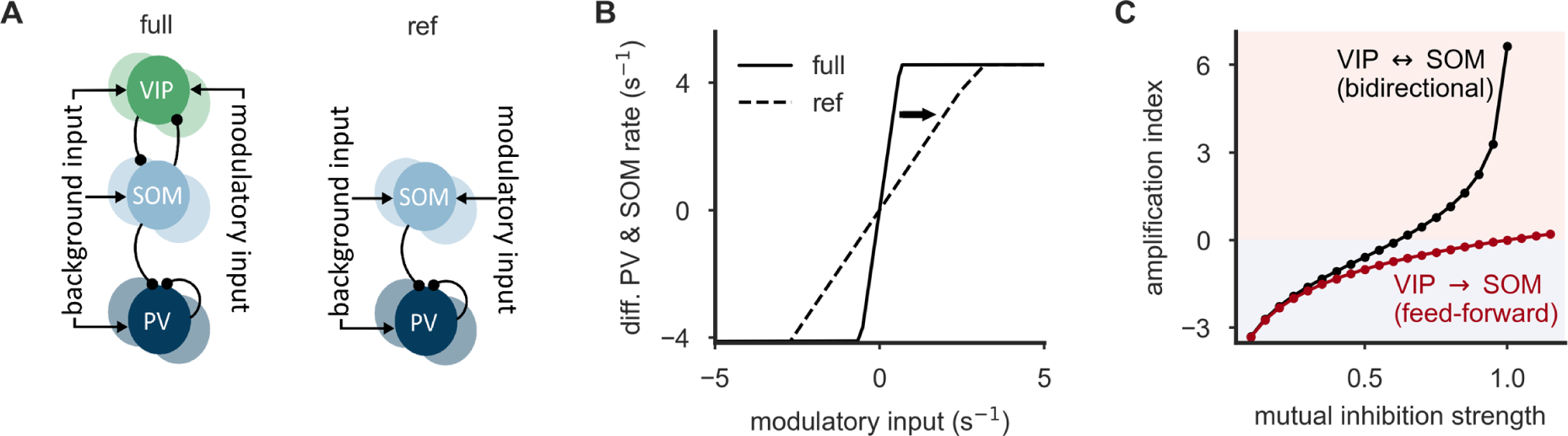
Mutual inhibition between SOM and VIP neurons enhances an amplification of weak input. **(A)** Reduced network of inhibitory PV, SOM and VIP neurons that allows a mathematical analysis, and a corresponding reference network without VIP neurons. **(B)** The difference of PV and SOM neuron rates (somatic and dendritic inhibition, respectively) follows a sigmoid function, the slope of which characterizes the gain of the redistribution of somato-dendritic inhibition upon a change in the modulatory input. The difference of PV and SOM neuron rates in the full interneuron network (A, left) exhibits a larger slope than a corresponding reference network without VIP neurons (A, right) (mutual inhibition strength *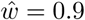*). **(C)** Amplification index depends strongly on the mutual inhibition strength. Positive values denote an amplification, negative values indicate an attenuation. An infinitely large amplification index corresponds to a winner-take-all (WTA) regime. When the connections from SOM to VIP neurons are knocked-out (red), the amplification index rises slowly and stays in the attenuation regime for a large range of total mutual inhibition strengths.

Again, we compared the circuit to a reference network without VIP neurons (Figure 2 A, right), in which modulatory inputs impinged directly onto the SOM neurons. In line with the full model, we observed that the removal of the VIP neurons led to a prominent reduction of the sensitivity to modulatory inputs, i.e., a reduced slope of the somato-dendritic difference of inhibition (cf. Figure 1 D and Figure 2 B). To quantify the effect of the SOM-VIP motif, we introduced an *amplification index A*, defined as the logarithm of the ratio of slopes in the two networks with and without VIP neurons (cf. Figure 2 B solid and dashed lines, and see Models and Methods for more details). An amplification index larger than zero indicates that the interneuron network amplifies weak input onto VIP neurons in comparison to the reference network.

The simplified circuit allows to derive a mathematical expression for the amplification index, which shows that the amount of amplification depends critically on two circuit properties (see Models and Methods for a detailed derivation). Firstly, it increases with the effective VIP→SOM connection strength, reflecting the monosynaptic effect of VIP inputs onto SOM neurons. Secondly, it depends in a highly nonlinear way on the product of the connection strengths from SOM*→*VIP and VIP*→*SOM. This second dependence is a consequence of the mutual inhibition between SOM and VIP neurons: an increase in VIP firing rate not only inhibits SOM neurons, but also further disinhibits the VIP neurons themselves, which in turn increase their rate, further inhibiting the SOM neurons etc. As this mutual inhibition approaches a critical strength, the amplification index increases rapidly. Beyond the critical strength, the circuit transitions into a competitive winner-take-all regime, in which either the SOM or the VIP neurons are silenced by the other population. Our mathematical analysis is confirmed by simulations, which also show a rapid increase of the amplification index as the mutual inhibition between VIP and SOM neurons increases (see Figure 2 C for symmetric mutual inhibition strengths, and Figure S1 for asymmetric weights). Much stronger VIP*→*SOM connection strengths are required to achieve an amplification (*A >* 0) when the back-projection SOM*→*VIP is knocked out, effectively eliminating the mutual competition between VIP and SOM neurons (red line in Figure 2 C). These results demonstrate that mutual inhibition is a key player in the amplification of weak inputs onto VIP cells.

### Connectivity and short-term plasticity support the amplification

If the SOM-VIP motif were to serve as an amplifier of weak modulatory signals, other circuit properties should also support this function. A candidate mechanism that would further enhance the competition between SOM and VIP is synaptic short-term faciliation (STF). Although short-term plasticity between different types of GABAergic interneurons has received limited attention, STF has indeed been demonstrated for the mutual connections between SOM and VIP neurons (Karnani et al., 2016). We therefore enhanced the network model by a Tsodyks-Markram type model of short-term plasticity (Tsodyks and Markram, 1997; Markram et al., 1998, see Models and Methods for more details). For the sake of simplicity, SOM VIP and VIP SOM synapses had equal facilitation parameters (Figure 3 A). The overall STF strength was varied by changing the initial release probability, while adjusting the synaptic weight in order to keep the initial postsynaptic response constant (see Models and Methods for further details). As expected, STF causes an increase of the amplification index, such that smaller mutual inhibition strengths are sufficient to achieve an amplification index above the amplification threshold *A* = 0 (Figure 3 A).

**Figure 3.**
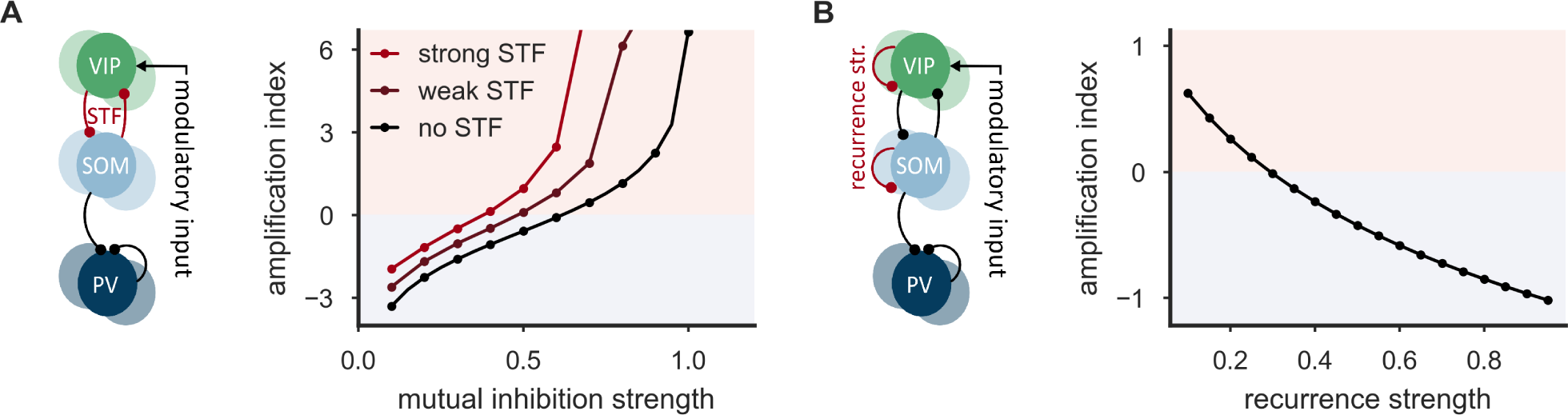
Connectivity properties of the SOM-VIP motif support the amplification in the interneuron network. **(A)** Network of inhibitory PV, SOM and VIP neurons with short-term facilitation (STF) of SOM*→*VIP and VIP*→*SOM connections (left). STF increases the amplification index. Smaller mutual inhibition strengths are sufficient to achieve an amplification index above the amplification threshold *A* = 0 (right). STF parameters are equal for SOM and VIP neurons: *U*_s_ = 0.1 (strong STF), *U*_s_ = 0.5 (weak STF), *U*_s_ = 1 (no STF) and *τ*_f_ = 100 ms. **(B)** Network of inhibitory PV, SOM and VIP neurons with artificially introduced recurrent connections among both SOM and VIP neurons (left). Recurrence leads to a decrease of the amplification index (right). Recurrence strengths are equal for SOM and VIP neurons. Mutual inhibition strength *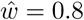*.

In contrast to PV neurons, which show strong inhibitory connections onto other PV cells (Pfeffer et al., 2013; Hu et al., 2014; Tremblay et al., 2016), SOM and VIP neurons only very rarely inhibit other neurons of the same class (Pfeffer et al., 2013; Jiang et al., 2015; Tremblay et al., 2016). To investigate whether this lack of recurrent inhibition supports the amplification properties of the network, we artificially introduced recurrent connections among both SOM and VIP neurons. We systematically varied their strength, while keeping the strength of mutual inhibition between the two populations constant. For simplicity, we considered a symmetric situation in which the strength of the recurrent inhibition is the same among VIP and SOM neurons (see Figure 3 B), but similar results are obtained in asymmetric situations (see Figure S2). We found that recurrent connections among SOM and VIP neurons lead to a strong reduction of the amplification index (Figure 3 B), even for relatively weak recurrent connections (see Models and Methods for a mathematical analysis). The strongest amplification was always observed for a connection strength of zero, that is, when recurrent inhibition is absent.

In summary, connectivity properties like short-term facilitation and the absence of recurrent connections among both VIP and SOM neurons support the effective translation of small stimuli onto VIP cells into large changes of somato-dendritic inhibition.

### Spike-frequency adaptation introduces a frequency-selective amplification

Both SOM and VIP neurons show an absence of recurrent inhibition within the same population, but they make use of a different negative feedback mechanism: spike-frequency adaptation (SFA). SFA is a prominent feature observed in many cortical neurons (La Camera et al., 2006; Tremblay et al., 2016). Upon stimulation, adapting cells decrease their firing rate gradually, and consequently exhibit a difference between steady-state and onset firing rate. SOM neurons feature salient SFA (Rudy et al., 2011; Gentet, 2012; Tremblay et al., 2016; Urban-Ciecko and Barth, 2016; Wamsley and Fishell, 2017), whereas VIP neurons claim a broad spectrum from weak to strong adaptation. To study the effect of SFA, we augmented the rate dynamics of SOM and VIP neurons by an additional rate adaptation variable (see Models and Methods). For simplicity, we again assumed the same adaptation parameters for SOM and VIP neurons.

While we expected both adaptation and recurrent inhibition to weaken the amplification, they differ with respect to the time scales on which they operate. Recurrent inhibition acts on the rapid time scale of synaptic transmission (e.g, of 5 - 10 ms for GABA_A_ receptor-based transmission). In contrast, adaptation operates on a wide range of time scales from tens to thousands of milliseconds (La Camera et al., 2006; Lundstrom et al., 2008; Pozzorini et al., 2013). Consequently, adaptation allows a gradual transition over time from amplification to attenuation.

To characterize the time dependence of the amplification properties, we performed a frequency response analysis (Figure 4 A-B), by stimulating VIP neurons with oscillating inputs of varying frequency. The difference of the population firing rates of PV and SOM neurons – as a reflection of the somato-dendritic distribution of inhibition – oscillates in response to this stimulation. The magnitude of this output oscillation, normalized by the amplitude of the input oscillation yields a frequency-resolved measure of amplification. We found that increasing recurrent inhibition among SOM and VIP neurons systematically reduces the output amplitude across all frequencies (Figure 4 A), confirming the steady-state analysis (cf. Figure 3 B). In contrast, spike-frequency adaptation introduces a prominent frequency selectivity: For low-frequency oscillations, the output amplitude decreases with increasing adaptation strength. For high-frequency oscillations, it increases (Figure 4 B-C). Furthermore, the circuit exhibits a preferred frequency (resonance frequency), for which it yields a maximal response. Neuronal adaptation hence introduces a frequency-selective amplification that preferentially transmits specific neuronal rhythms within the broad spectrum of oscillations in the brain (Buzsáki and Draguhn, 2004).

**Figure 4.**
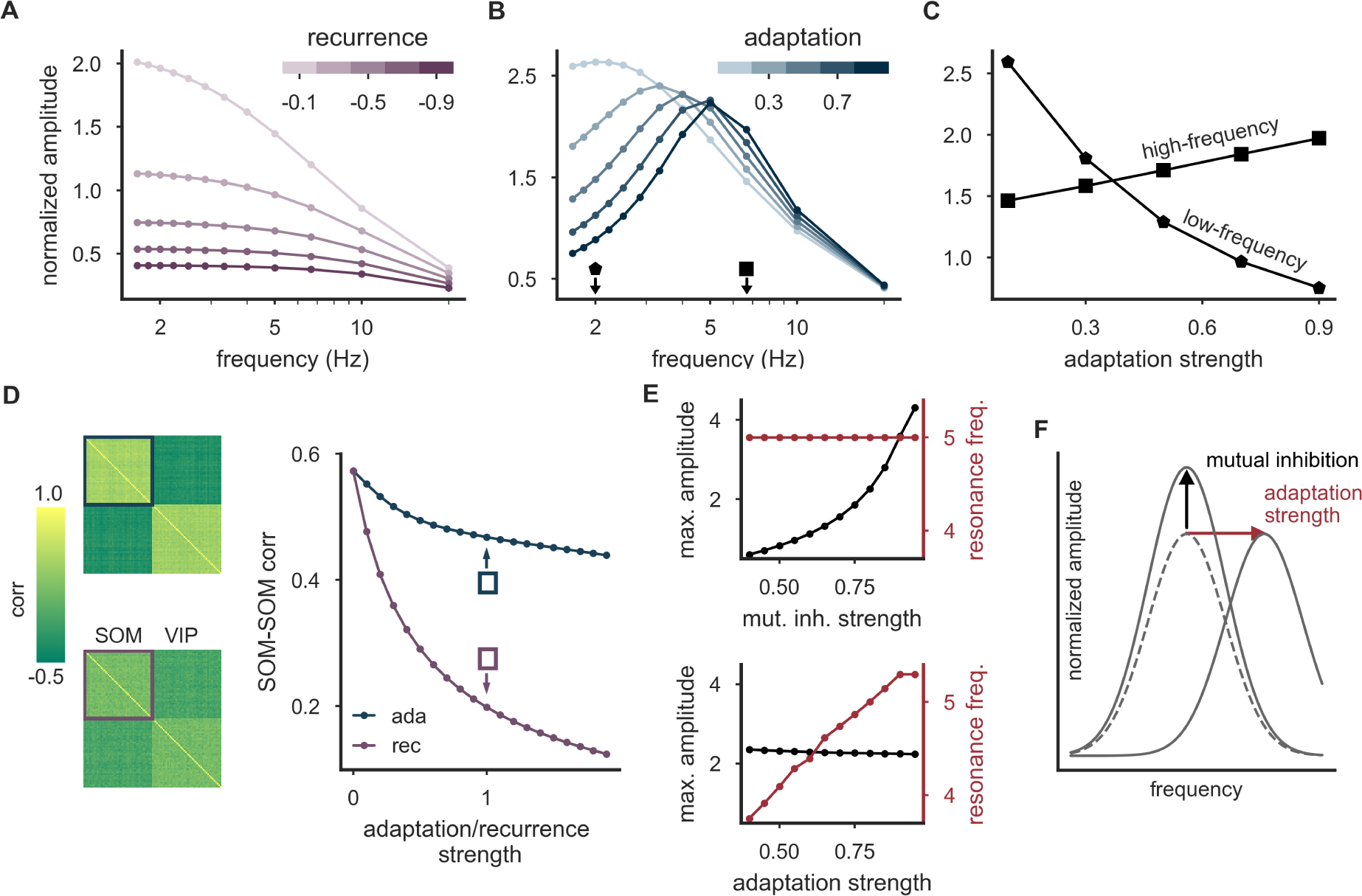
spike-frequency adaptation enables frequency-selective amplification and preserves co-activity. **(A)** Frequency response analysis of the interneuron network with recurrent connections among both SOM and VIP neurons. Increasing recurrence reduces the oscillation amplitudes across all stimulation frequencies. **(B)** Same as in A, but with spike-frequency adaptation instead of recurrence. The circuit yields a maximal response at a resonance frequency. With increasing adaptation strength, this resonance frequency increases. **(C)** The normalized amplitude decreases with increasing adaptation strength for low-frequency oscillations, but increases for high-frequency oscillations (frequency-selective amplification, cf. markers in (B)). **(D)** Neurons show stronger correlations (positive or negative) with spike-frequency adaptation (left, upper panel) than with recurrent inhibition among SOM and VIP neurons (left, lower panel). Mean SOM-SOM neuron correlation is more sensitive to increasing recurrence strength than to adaptation strength (right). **(E)** While strengthening mutual inhibition leads to an increase of the maximal frequency-resolved measure of amplification, it does not change the resonance frequency (top, adaptation strength *b* = 0.8). In contrast, increasing the adaptation strength allows to adjust the resonance frequency, with a weak impact on the maximum of the oscillation amplitude (bottom, recurrence strength *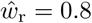*). **(F)** Schematic representation of both separate “knobs” (mutual inhibition and adaptation) and their independent control of the amount and frequency-selectivity of the amplification. Parameter (A-E): Mutual inhibition strength *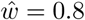*.

Spike-frequency adaptation and recurrent inhibition also have distinguishable consequences for the correlation structure of the interneuron network. Karnani et al. (2016) demonstrated that both SOM and VIP neurons are cooperatively active as populations rather than individually. We studied this co-activity by stimulating both interneuron populations with shared and individual noise on top of a constant background input. The shared noise between members of the same interneuron class introduced strong correlations between both VIP/VIP and SOM/SOM neurons as described by Karnani et al. (2016). We then studied how recurrent inhibition and adaptation differentially affect the co-activity of the populations, quantified by the averaged pairwise correlation coefficient. We found that recurrent inhibition strongly decreases the correlation between members of the same interneuron class (Figure 4 D; Renart et al., 2010), while adaptation introduces only a marginal reduction, and thereby preserves the high correlations seen by Karnani et al. (2016).

As for any amplifier, it would be useful to allow a dynamic adjustment of the amplification in the circuit. Very strong amplification would lead to small ranges of effective modulatory signals and a rapid saturation of the system, and hence to potential distortions in the translation of the modulatory signal into somatic and dendritic inhibition. Similarly, it could be beneficial to allow an adjustable frequency-based selection of the modulatory input. A neuronal mechanism that is suitable to tune these two properties could be a neuromodulatory control of circuit parameters (Hasselmo, 1995). In simulations, we found that a strengthening of mutual inhibition increases the overall amplification, while leaving the resonance frequency largely unaltered (Figure 4 E, top). At the same time, changes of the adaptation strength allow to tune the resonance frequency, while leaving the maximum of the frequency-resolved measure of amplification largely unchanged (Figure 4 E, bottom). In summary, the circuit seems to display separate “knobs”, which offer an independent control of the amount and frequency-selectivity of the amplification through separate neuromodulatory channels (Figure 4 F).

These results demonstrate that spike-frequency adaptation, though similar in its steady-state properties to recurrent inhibition within SOM and VIP populations, enables a frequency-selective amplification with well-separated target parameters for neuromodulatory control.

### The computational repertoire of the SOM-VIP motif

Our simulations indicate that the SOM-VIP subnetwork supports different computational functions, ranging from signal amplification and frequency selection to switching behavior for strong mutual inhibition between the populations. To understand how these computational states are determined by the parameters of the system, we ran extensive simulations of the SOM-VIP motif alone, accompanied by mathematical analyses of a linearized network without rate rectification.

We first investigated a network of VIP and SOM neurons that is consistent with experimentally observed characteristics, i.e., with adaptation and inhibitory connections exclusively between neurons of different type. Simulations reveal four operating modes for such a network (Figure 5 A). For weak mutual inhibition, the two interneuron populations can be active at the same time, and modulatory VIP signals are attenuated (Figure 5 A, region a). As the inhibition between the two populations increases, the amplification index increases and leads the circuit into an amplification domain (Figure 5 A, region b). Beyond a critical strength of mutual inhibition, the network then transitions into a switch-like winner-take-all domain (Figure 5 A, regions c & d), in which one of the two populations silences the other. Notably, this state comes in two variants: For weak adaptation, the winning population silences the other permanently, or until an external event switches the network to a different winner (Figure 5 A, region c). We will show later that this allows transient VIP inputs to persistently switch the operating mode in the full microcircuit. For strong adaptation, the network shows oscillatory switching (Figure 5 A, region d), because adaptation gradually decreases the firing rate of the winning interneuron population. This releases the other population from inhibition until it can no longer be silenced and becomes the new winner and in turn starts to adapt. A mathematical analysis of the linearized network predicts the parameter ranges of the four computational states almost perfectly (see black lines in Figure 5 A, and Models and Methods for more details). The observed oscillations comprise a wide spectrum of frequencies that depend non-linearly on the strength and time constant of adaptation and on the strength of mutual inhibition (Figure 5 B, see also Figure S3 and S4 for networks with asymmetric adaptation parameters for SOM and VIP neurons). Deviations between the frequencies observed in simulations and those predicted by the mathematical theory are caused by the omission of the rate rectification in the theory. The four computational states of the network are also observed when short-term plasticity is introduced into the network, although the transition boundaries change such that the switch-like state is reached for weaker mutual inhibition and the oscillatory switch requires stronger adaptation (Figure 5 C).

**Figure 5.**
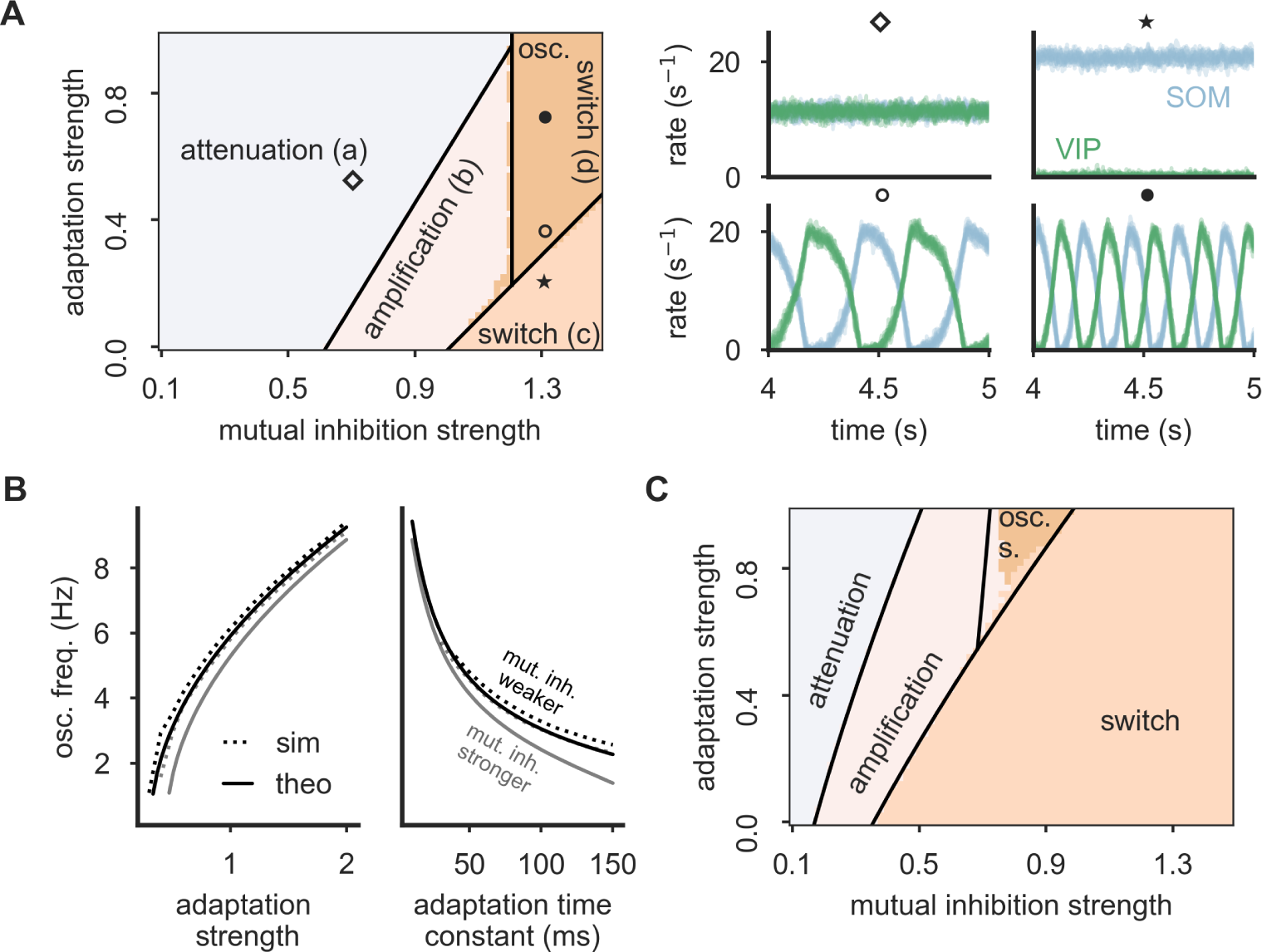
Dynamical states of the SOM-VIP motif. **(A)** Left: Bifurcation diagram reveals distinct operation modes: all interneurons are active (divided into amplification (a) and attenuation regime (b)), a winner-take-all (WTA) regime implementing a switch (c) and an oscillatory WTA regime leading to an oscillating switch (d). Regime boundaries (black lines) are obtained from a mathematical analysis (see Appendix). Right: Example firing rate traces for all SOM (blue) and VIP (green) neurons for four network settings taken from the bifurcation diagram (cf. markers). Adaptation time constant *τ*_a_ = 50 ms. **(B)** The oscillation frequency in the oscillatory switch mode depends on the adaptation strength (left), the adaptation time constant (right) and the total mutual inhibition strength (black: *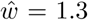* = 1.3, gray: *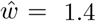* = 1.4). Left: *τ*_a_ = 50 ms, right: *b* = 1. The frequencies cover a broad range from Delta to Alpha oscillations. **(C)** When short-term facilitation (STF) is present, the WTA (switch) regime is enlarged and the switch-like oscillation mode requires stronger adaptation. Initial synaptic efficacy *U*_s_ = 0.1, facilitation time constant *τ*_f_ = 100 ms.

Two inhibitory populations that mutually inhibit each other may well be a common network motif in cortical circuits, and the absence of recurrent inhibitory connections within the two populations – as observed in the SOM-VIP motif – may not always hold.

We therefore also performed an analysis of the computational states of a network with recurrent inhibition. Simulations reveal five operating modes for such networks (see Figure S5). When mutual inhibition and recurrent inhibition is weak, we again found that both interneuron populations can be active at the same time. Depending on the strength of the mutual inhibition, we again observed attenuation and amplification, respectively. For sufficiently strong mutual inhibition between SOM and VIP cells, the amplification regime transitions into the switch-like state where only one population is active. In contrast to adapting neurons, the network did not show an oscillatory state. Instead, very strong recurrent inhibition introduces strong competition between the neurons within the interneuron populations, leading to pathological states where either one single cell per cell type is active (if mutual inhibition is weak) or only one single neuron at all is active (if mutual inhibition is strong). Again, these dynamical states and their transitions are predicted almost perfectly by a mathematical eigenvalue analysis (see black lines in Figure S5, and Models and Methods for derivation). The mathematical analysis also unveils that for sufficiently large populations, the pathological states require very strong synapses (ultimately, a single cell must silence all others) and are hence unlikely to be observed in the nervous system.

In summary, the SOM-VIP network motif allows different computational states, covering attenuation, amplification, switching and – for adapting neurons – oscillatory switching in a frequency range of Delta (1-4 Hz), Theta (4-8 Hz) or Alpha (8-12 Hz) oscillations.

### Switch between distinct processing modes in local microcircuits

To investigate the computational consequences of the SOM-VIP circuit, we returned to the full microcircuit comprising PCs and inhibitory PV, SOM and VIP cells (Figure 6 A left). We first verified that all results observed in the simplified interneuron networks still hold for the larger circuit. Again, stronger mutual inhibition and the presence of STF increase, while negative feedback mechanisms like recurrent inhibition or adaptation decrease the system’s sensitivity the modulatory input (Figure 6 A right). Furthermore, adaptation leads to a frequency-selective amplification for which the processing of rapidly changing input signals benefits from powerful spike-frequency adaptation in SOM and VIP neurons (Figure 6 B). Finally, the experimentally observed elevated correlation between members of the same interneuron class is also preserved for adaptation and decreases strongly for recurrent inhibition (Figure 6 C). In summary, the results obtained in the simplified interneuron network also hold in the full circuit.

**Figure 6.**
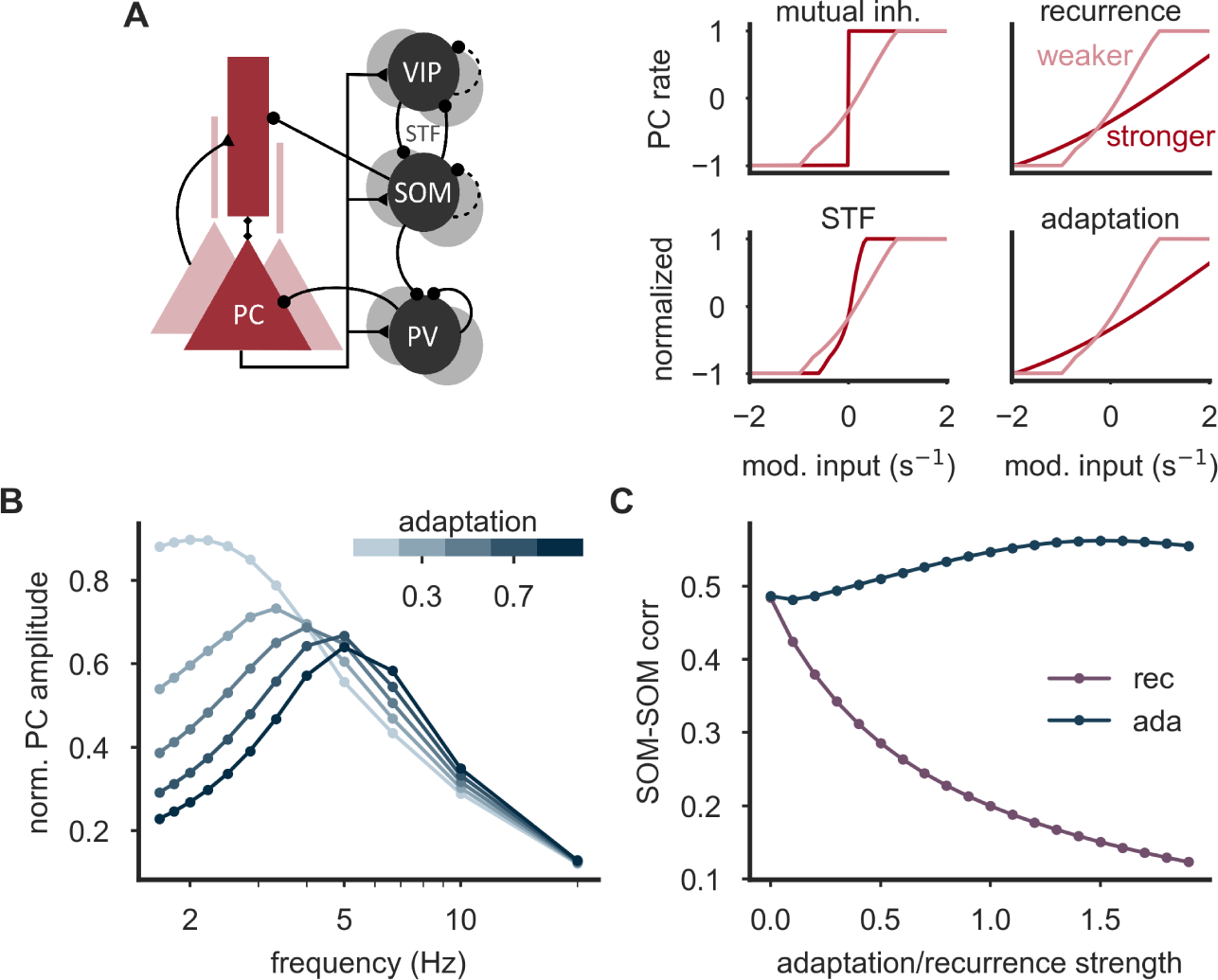
The full microcircuit with excitatory PC and inhibitory PV, SOM and VIP neurons shows the same phenomena as the reduced interneuron network. **(A)** PCs in the full microcircuit (left) exhibit a steeper firing rate slope with increasing mutual inhibition strength (*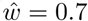* = 0.7 and 0.9), increasing short-term facilitation (STF) (*U*_s_ = 0.4 and 0.25, *τ*_f_ = 200 ms), but a reduced slope for increasing recurrence strength (*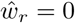*_*r*_ = 0 and 0.3) and adaptation (*b* = 0.2 and 0.5, *τ*_*a*_ = 100 ms). **(B)** With adaptation, the frequency response analysis reveals a frequency-selective amplification. **(C)** Mean SOM-SOM neuron correlation is more sensitive to increasing recurrence strength than to adaptation strength (right). Parameter (B & C): Mutual inhibition strength *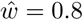*.

What is the computational impact of a somato-dendritic redistribution of inhibition on PCs? It is well established that on their apical dendrites, many pyramidal cells receive top-down input from higher cortical areas (Felleman and Van, 1991; Cauller et al., 1998) and matrix thalamic nuclei (Diamond, 1995). On their basal dendrites and the perisomatic domain, they receive bottom-up input from lower cortical areas and core thalamic nuclei (Larkum, 2013). Although inputs at the electrically distant apical dendrites have a small impact on initiating spikes at the axon initial segment (Stuart and Spruston, 1998; Williams and Stuart, 2002), they can initiate long-lasting calcium spikes when they coincide with back-propagating action potentials from the soma (Yuste et al., 1994; Larkum et al., 1999b; Larkum and Zhu, 2002), leading to a significant gain increase of L5 pyramidal cells (Larkum et al., 2004). How the two streams of information are integrated is not fully resolved (Naud and Sprekeler, 2018). In particular, it is conceivable that top-down inputs are only used when they provide useful and reliable information, and are ignored otherwise. Given that calcium spikes in apical dendrites are very sensitive to inhibition (Larkum et al., 1999a), dendrite-targeting interneurons are well suited to control the integration of top-down inputs, and, consequently, the switch between distinct modes of operation. We therefore simulated the full microcircuit with top-down and bottom-up inputs, in a “switch” configuration of strong mutual inhibition between VIP and SOM, and without spike-frequency adaptation. For illustration, we chose as inputs two sinusoidal oscillations with different frequencies (Figure 7 A & C), and determined how strongly these two input streams are represented in the firing rate of PCs (Figure 7 A & B). When the network is in the “switch” regime, we found that small changes in VIP input are sufficient to switch the network between two computational states in which top-down inputs are either transmitted or cancelled entirely (Figure 7 B). Interestingly, transitions between those two computational states can also be triggered by transient modulatory pulses. The evoked change of the processing mode persists even after the end of the pulse (Figure 7 C), reflecting a bistability of the system. As a consequence, the current processing mode of the network depends on the recent history of the input to the SOM-VIP motif (Figure S6).

**Figure 7.**
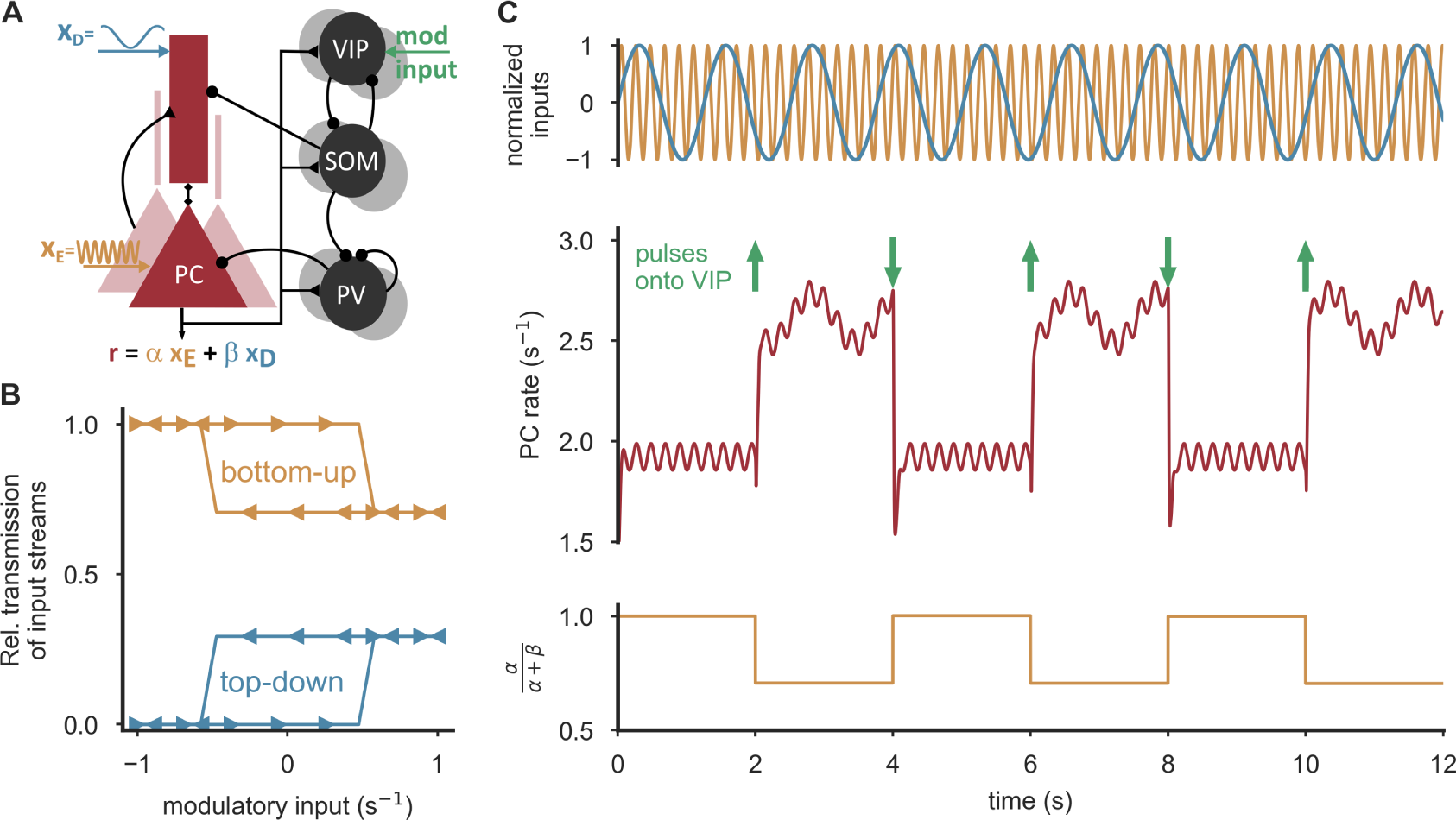
Integration or cancelation of top-down signals by modulatory VIP input. **(A)** PCs of the full microcircuit are stimulated at the soma and the apical dendrites with two oscillations with different frequencies, emulating bottom-up (orange, *x*_E_) and top-down (blue, *x*_D_) input, respectively. PC rate reflects the two inputs *x*_E_ and *x*_D_ with different coefficients *α* and *β*, depending on the modulatory input. **(B)** In an amplification regime (*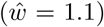* = 1.1), weak, permanent modulatory VIP input is sufficient to switch between two operation modes, in which top-down input is either integrated (*β >* 0) or canceled (*β* = 0). In the WTA regime, the network exhibits hysteresis, that is, the level of modulatory input needed to cause a switch depends on the network state. For a range of inputs, the circuit is bistable. **(C)** In the bistable regime (*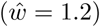* = 1.2), persisting transitions between the states can be triggered by strong, short pulses delivered to VIP neurons (10 ms duration, amplitude 8.4/s, timing denoted by green arrows). Parameters (A-C): Weight between SOM neurons and dendrites *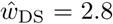*_DS_ = 2.8, External stimulation x_E_ = 25*/*s + 0.5 sin(5 *t*)*/*s, x_D_ = 7*/*s + 0.1 sin(30 *t*)*/*s, x_PV_ = 12/s, x_SOM_ = x_VIP_ = 3.5/s.

In summary, we demonstrate that the integration of top-down feedback from higher cortical areas can be induced or prevented by persistent, weak input or short, strong input pulses onto VIP cells. As the network exhibits hysteresis, the switching depends on the collective state of SOM and VIP neurons.

### Amplification of small mismatch signals

The switching property of the circuit relies on a competition between VIP and SOM neurons. Hence, which of the two populations dominates should not be determined by the input to VIP neurons alone, but rather by the (potentially weighted) difference between the inputs to SOM and VIP neurons (Wong and Wang, 2006). By systematically varying both inputs in an amplification regime, we indeed found that the somato-dendritic distribution of inhibition is determined by the difference between the two inputs (Figure 8 A). The SOM-VIP circuit can hence be interpreted as an amplifier for small differences between two input streams that impinge onto SOM and VIP neurons.

**Figure 8.**
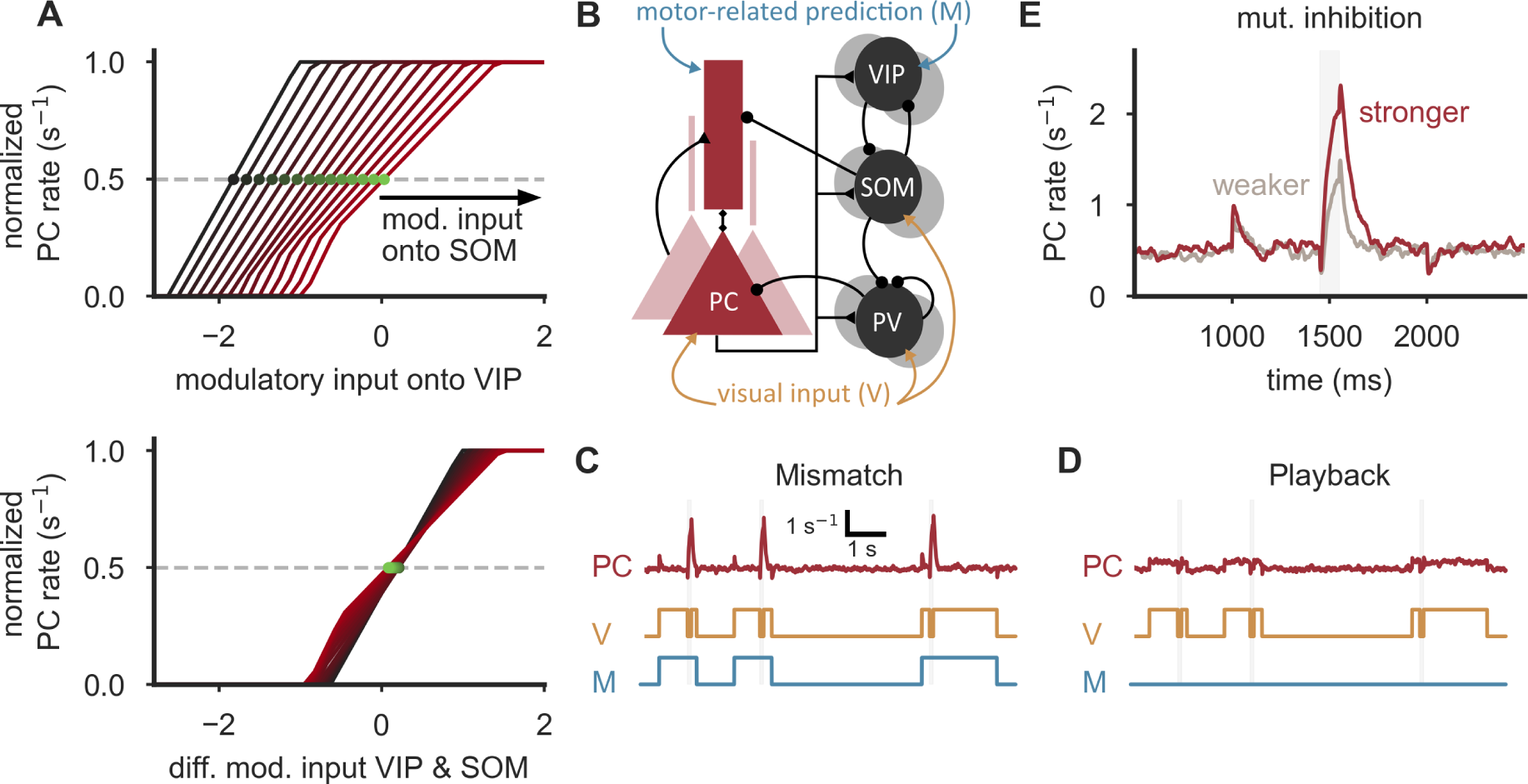
Mismatch detection by SOM/VIP-dependent amplification. **(A)** Additional modulatory input onto SOM neurons shifts the transition point (green) of the PC rate as a function of modulatory VIP input (top). When plotted as a function of the difference of SOM and VIP input, the transition points align, indicating that the circuit amplifies differences between two input streams (bottom). Additional input onto SOM neurons ranges from −2/s to 0/s. **(B)** Model for the integration of visual inputs and motor-related predictions (Attinger et al., 2017). SOM neurons, PV neurons and the somatic compartment of PCs receive external visual input, VIP neurons and the apical dendrites of PCs receive an internal (motor-related) prediction of the expected visual input. The connection strengths from PV neurons to the somatic compartment of PCs and the SOM → PV connection were chosen to ensure a response only when the visual input is switched off and the (motor-related) prediction is switched on (see Methods). **(C)** PC neurons respond with an increase in firing rate when visual input is off and the motor-related input is on (mismatch), but show a negligible increase in activity above baseline when both input streams are on. **(D)** Also, only negligible responses above baseline are evoked when motor-related input is permanently off (playback session). Mutual inhibition strength *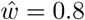*. **(E)** The mismatch-induced increase in firing rate is more pronounced in an amplification regime (*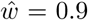*, dark red) in comparison to an attenuation regime (*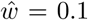*, gray). Parameters (B-E): Motor-related and visual input on corresponds to an additional input of 10.5/s and noise drawn from a Gaussian distribution with zero mean and SD = 3.5/s. Background stimulation x_E_ = 24.5/s, x_D_ = 0/s. Time constant of PC neurons increased by factor 6 to reduce onset responses.

This observation is interesting in the context of a recent study of Attinger et al. (2017). The authors suggested a conceptual model for layer 2/3 of mouse V1, in which SOM neurons receive visual inputs, while VIP neurons and the apical dendrites of the PCs receive an internal (motor-related) prediction of the *expected* visual input. When properly tuned, the excitatory top-down input to the PC dendrites is then cancelled by SOM inhibition, as long as the internal prediction matches the sensory data. Deviations between sensory inputs and internal predictions, however, change the level of dendritic inhibition and thereby generate mismatch responses, as observed in a subset of PCs in V1 (Keller et al., 2012; Attinger et al., 2017) and in other systems (Keller and Hahnloser, 2009; Eliades and Wang, 2008).

To test this hypothesis *in silico*, we stimulated our full circuit model with visual inputs – impinging onto PV and SOM neurons and the somata of PCs – and motor feedback – impinging onto VIP neurons and the apical compartment of PCs (Figure 8 B). Similar to the findings of Keller et al. (2012), we found network configurations in which we observed selective responses in PCs when motor feedback was present in the absence of visual stimulation (Figure 8 C), but not when visual stimuli were presented in the absence of motor feedback (Figure 8 D). These responses were reduced when the circuit was brought into an attenuation configuration by weakening mutual inhibition between VIP and SOM neurons (Figure 8 E).

We therefore suggest that the amplification brought about by the competition between

SOM and VIP neurons could serve to amplify small deviations between different input streams, such as sensory signals and internal predictions. Because such deviations are powerful learning signals for the internal prediction system (Wolpert et al., 2011), an amplification may be beneficial for learning highly accurate predictions.

## Discussion

We have shown that the broadly observed microcircuit comprising excitatory PC and inhibitory PV, SOM and VIP neurons can act as an amplifier that translates weak input onto VIP cells into large changes in the somato-dendritic distribution of inhibition onto PCs. A cornerstone of this amplification is mutual inhibition between SOM and VIP neurons that - if sufficiently strong - allows switch-like spatial shifts of somato-dendritic inhibition. Connectivity properties like short-term facilitation of those mutual connections and the absence of recurrent connections among both SOM and VIP neurons support the amplification. Spike-frequency adaptation as observed for SOM and VIP cells gives rise to a frequency-selective amplification, as a consequence of the slow time scales of adaptation. Furthermore, adaptation in conjunction with sufficiently strong mutual inhibition results in an oscillatory switching regime (Laing and Chow, 2002; Moreno-Bote et al., 2007), in which SOM and VIP neurons alternately win the competition, thus generating rhythmic shifting of somato-dendritic inhibition. These oscillations are inherited by PC neurons that fluctuate between two different computational states with frequencies ranging from approximately 1 to 10 Hz. Functionally, such oscillations could reflect a rhythmic switching between a state in which sensory data is acquired and a state in which it is calibrated against internal predictions.

### Evidence for model assumptions

The microcircuit we studied has been observed in several cortical areas, including mouse primary somatosensory (S1), visual (V1) and vibrissal motor (vM1) cortex, and both in layer 2/3 and 5 (Bartley et al., 2008; Fino and Yuste, 2011; Packer and Yuste, 2011; Avermann et al., 2012; Pfeffer et al., 2013; Lee et al., 2013; Jiang et al., 2015; Jouhanneau et al., 2015; Pala and Petersen, 2015; Urban-Ciecko et al., 2015). We did not strive to resolve subtle differences between these areas, but rather covered broad parameter ranges to explore the computational repertoire of the circuit (see, e.g., Wang and Yang, 2018, for area-to-area variations). Whether the SOM-VIP motif would act as an amplifier, switch or even a rhythmic switch in these different regions will depend on details of the circuit. In the following, we discuss a few assumptions made in the model, including the absence of recurrent inhibition within the SOM and VIP populations and the fact that we ignored VIP connections onto other cell classes.

The absence of recurrent connections among both SOM and VIP neurons has been supported by many studies, both in vitro and in vivo (e.g., Pfeffer et al., 2013; Karnani et al., 2016). It has been argued, however, that SOM neurons should be subdivided into ‘Martinotti’ and ‘non-Martinotti’ cells to account for subtle differences in their morphology and biophysical properties (e.g., Jiang et al., 2015). While recurrence seems to be weak or absent within both sub-populations, connections between these subgroups have been reported. However, most of the SOM neurons belong to the class of Martinotti cells that avoid connections to each other. Moreover, the number of ‘non-Martinotti’ cells, their connection probability and strength is relatively weak, such that our assumption of no recurrence between SOM neurons is a reasonable approximation (Jiang et al., 2015).

The model contained a unidirectional connection from SOM neurons onto PV neurons. This assumption is based on the common observation that this connection is strong and frequent, while the backprojection is rather weak or absent in layer 2/3 and 5 (see Pfeffer et al., 2013; Urban-Ciecko and Barth, 2016). An exception is the study of Walker et al. (2016), who reported strong and frequent connections from PV to Martinotti cells in layer 2/3 of mouse S1. We expect that such a mutual inhibition between PV and SOM neurons would not violate our hypothesis, but rather introduce another amplification mechanism with a similar structure. However, PV → SOM connections may also introduce complex interactions between inputs to PV and VIP neurons that are not captured in our model. It remains for future work to explore the computational and functional consequences of such serially connected amplifiers.

In our model, VIP neurons are connected exclusively to SOM neurons. We neglected potential VIP input to PCs and other interneuron classes. This assumption is supported by a wealth of experimental studies that reported no or weak connections from VIP to PC and PV neurons (e.g. see Pi et al., 2013; Pfeffer et al., 2013; Jiang et al., 2015), and a net disinhibitory impact of VIP neurons onto PCs in S1 (Lee et al., 2013), V1 (Pfeffer et al., 2013; Pi et al., 2013; Fu et al., 2014; Zhang et al., 2014) and the auditory cortex (Pi et al., 2013). However, in a study of Garcia-Junco-Clemente et al. (2017) strong and direct connections between VIP and PC neurons were found during arousal in layer 2/3 of the mouse frontal association area. The strength of this inhibition was highly variable between cells, covering a wide range of almost two orders of magnitude. Also, this connection was reported to be weaker in the occipital cortex in the same study (Garcia-Junco-Clemente et al., 2017). Notwithstanding the presence of this connection in different systems, it is tempting to speculate on its computational function, because it may enable to run different operating modes in parallel. Karnani et al. (2016) have demonstrated that SOM and VIP neurons receive local excitation from distinct (non-overlapping) PC groups rather than non-selectively from all nearby excitatory neurons. If the respective interneuron types also selectively project back, and if the VIP projection would also impinge onto apical dendrites, one group of PC neurons may operate in a feedback-modulated state, whereas another group runs in a feedforward-driven mode at the same time (see Figure 9).

**Figure 9.**
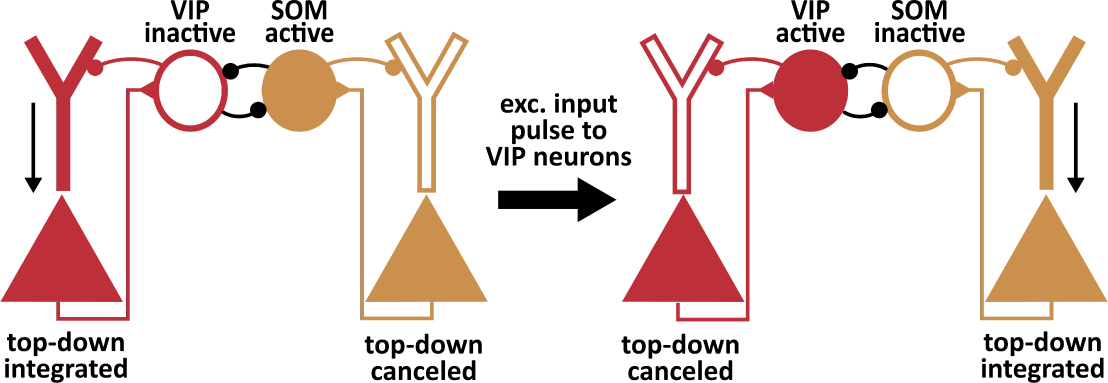
Diverse processing modes at the network level. Garcia-Junco-Clemente et al. (2017) suggested that VIP neurons also target the apical dendrites of PCs. Moreover, Karnani et al. (2016)) suggested that SOM and VIP neuron populations receive excitatory input from distinct (non-overlapping) PC populations. If SOM and VIP interneurons preferably inhibited those PCs from which they get their input, it is conceivable that the distinct PC populations operate in a feedback-modulated state and a feedforward-driven mode, respectively. The switch between these states could then be triggered by VIP neuron input or the difference between modulatory input onto SOM and VIP cells.

The modulatory VIP neuron input in our microcircuit represents an abstract control signal. We deliberately did not specify the origin of this signal throughout most of this study, because inputs to VIP cells are very diverse. Besides local excitation from PC neurons in the same and deep layers, the greatest source of excitatory input of VIP cells is feedback from higher cortical areas and thalamus (Harris and Shepherd, 2015; Tremblay et al., 2016; Wall et al., 2016). Moreover, VIP neurons are also strongly excited by acetylcholine and serotonin (Harris and Shepherd, 2015), and Pi et al. (2013) have shown that VIP neurons in the auditory and prefrontal areas are recruited by reinforcement signals during an auditory discrimination task. Furthermore, in barrel cortex, VIP cells increase their activity during whisking as they receive substantial input from vM1 pyramidal neurons (Lee et al., 2013). Finally, it has been shown that locomotion activates VIP cells in V1 (independent of visual stimulation) and their firing rate is correlated with running speed (Fu et al., 2014). In summary, the assumption of a modulatory signal targeting VIP cells is supported by experimental data, but its origin or functional meaning may well vary between areas or over time.

### Testing the hypothesis

The hypothesis that the SOM-VIP motif mediates an amplification remains to be tested in experiments. The most direct test would be a measurement of the amplification index with VIP neurons active or deactivated. This could be achieved, e.g., by comparing the population activity of SOM neurons before and after silencing VIP cells optogenetically for a range of inhibitory and excitatory inputs presented to VIP and SOM neurons, respectively.

A central prerequisite of the hypothesis is that mutual inhibition between SOM and VIP neurons is strong. What does *strong* inhibition mean here? How does the unitless strength of synaptic connections in the model translate into physiological quantities? An intuition can be obtained from the amplification mechanism. The transition into switch-like behavior occurs when the total disynaptic disinhibition in the SOM-VIP motif is stronger than existing negative feedback mechanisms, such as leak or adaptation currents (or recurrent inhibition, if present). Consequently, the total inhibitory conductance a cell receives from the other interneuron type needs to be comparable to its own leak, a hallmark that could be observed physiologically.

### Functional implications

The ability to switch between distinct operating modes increases the computational repertoire of principal cells. When SOM neurons are inactive, top-down inputs onto apical dendrites elevate the output of pyramidal cells, for instance by bursts generated by dendritic calcium spikes (Larkum, 2013). On the other hand, when SOM neurons are highly active, the transmission of dendritic signals can be effectively canceled. Feedback “top-down” projections have been associated with a variety of cognitive parameters, including attention and visual awareness (Lamme et al., 1998; Siegel et al., 2000; Pascual-Leone and Walsh, 2001; Spratling, 2002), context (Tomita et al., 1999; Siegel et al., 2000; Olson et al., 2001; Spratling, 2002), internal predictions of the outside world (Rao and Ballard, 1999; Siegel et al., 2000; Larkum, 2013) and error or reinforcement signals for learning (Spratling, 2002; Rumelhart et al., 1986; Guerguiev et al., 2017; Sacramento et al., 2017). Hence, the integration or cancellation of top-down inputs from higher to lower cortical areas is likely to play a crucial role in information processing, cognition and perception (Siegel et al., 2000; Pascual-Leone and Walsh, 2001; Larkum, 2013). The SOM-VIP microcircuit could enable an efficient control over these processes by cortical and thalamic inputs to VIP neurons (Lee et al., 2013; Fu et al., 2014; Zhang et al., 2014; Wall et al., 2016).

Modulatory inputs may not comprise inputs to VIP neurons alone. The ability to amplify weak signals arriving in the SOM-VIP motif may be of particular importance in the context of detecting mismatches between two sources of information (Keller et al., 2012; Attinger et al., 2017). This could allow not only to amplify, but also to compute error signals that drive the refinement of internal models or the computational function of hierarchical “deep” networks (Spratling, 2002; Rumelhart et al., 1986; Guerguiev et al., 2017; Sacramento et al., 2017).

Despite the accumulating data on the broad variety of interneurons (Tremblay et al., 2016), their computational function is still poorly understood. The present study provides a hypothesis for one candidate role, which may well be only one in a broad repertoire of functions performed in parallel. Computational models may offer a useful resource to understanding this functional repertoire (Hayut et al., 2011; Litwin-Kumar et al., 2016; Yang et al., 2016; El-Boustani and Sur, 2014; Lee and Mihalas, 2017), given that they offer a degree of control over the circuit that is hard to achieve in experimental setups.

## Models and methods

### Neural network model

We simulated a rate-based network of excitatory pyramidal cells (*N*_PC_ = 70) and inhibitory PV, SOM and VIP cells (*N*_PV_ = *N*_SOM_ = *N*_VIP_ = 10). All neurons are randomly connected with connection probabilities (see Table 1) consistent with the experimental literature (Avermann et al., 2012; Pfeffer et al., 2013; Jiang et al., 2015). If not stated otherwise, all cells of the same neuron type have the same number of incoming and outgoing connections, respectively. This assumption is made merely for purposes of mathematical tractability and does not qualitatively alter the results.

**Table 1.** Connection probabilities between neuron types. Entries in the same columns correspond to the same presynaptic neuron type, entries in the same row to the same postsynaptic neuron type. Parentheses denote values that are only used when recurrence is introduced artificially. E: somatic PC compartment, D: dendritic PC compartment.

|  | E (PC som.) | PV | SOM | VIP |
| --- | --- | --- | --- | --- |
| E (PC som.) | - | 0.6 | - | - |
| D (PC dendr.) | 0.1 | - | 0.55 | - |
| PV | 0.45 | 0.5 | 0.6 | - |
| SOM | 0.35 | - | (0.5) | 0.5 |
| VIP | 0.1 | - | 0.45 | (0.5) |

The excitatory pyramidal cells are simulated by a two-compartment rate model taken from Murayama et al. (2009). The steady-state firing rate *r*_E,*i*_ of the somatic compartment of neuron *i* obeys

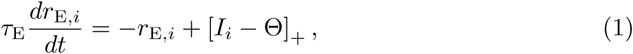

where [*x*]_+_ = *max*(0, *x*) is a rectifying nonlinearity and *τ*_E_ denotes a rate time constant (*τ*_E_=10 ms, unless stated otherwise). Θ denotes the rheobase of the neuron and *I*_*i*_ is the total somatic input generated by somatic and dendritic synaptic events and potential dendritic calcium spikes,

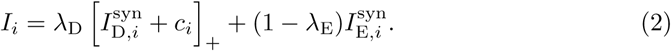

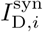 and 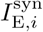 are the total synaptic inputs into dendrites and soma, respectively, and *c*_*i*_ denotes the dendritic calcium event. *λ*_D_ and *λ*_E_ are the fraction of “currents” leaking away from dendrites and soma, respectively. The synaptic input to the soma 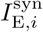 is given by the sum of external bottom-up in puts *x*_E_ and PV neuron-induced (P) inhibition,

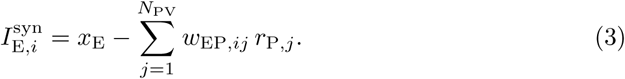

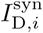 is the sum of top-down inputs *x*_D_, the recurrent, excitatory connections from other PCs and SOM neuron-induced (S) inhibition:

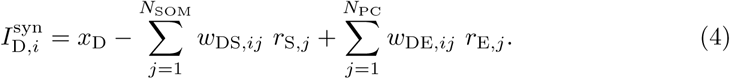

The weight matrices *w*_EP_, *w*_DS_ and *w*_DE_ denote the strength of connection between PV neurons and the soma of PCs (*w*_EP_), SOM neurons and the dendrites of PCs (*w*_DS_) and the recurrence strength between PCs (*w*_DE_), respectively (see Table 2). Note that all existing connections between neurons of type X and Y have the same strength, *w*_XY,*ij*_ = *w*_XY_, *X, Y*∈ {*E, D, P, S, V* } All weights are scaled in proportion to the number of existing connections (i.e., the product of the number of presynaptic neurons and the connection probability), so that the results are independent of population sizes.

The input generated by a calcium spike is given by

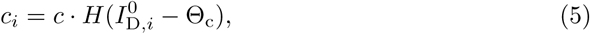

where *c* scales the amount of current, *H* is the Heaviside step function, Θ_c_ represents a threshold that describes the minimal input needed to produce a Ca^2+^-spike and 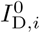 denotes the total, synaptically generated input in the dendrites,

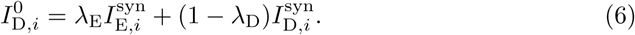

Unless stated otherwise, parameters were taken from Murayama et al. (2009) (see Table 2). Note that we incorporated the gain factor present in Murayama et al. (2009) into the parameters to achieve unit consistency for all neuron types.

**Table 2.** PC parameters describing the two-compartment rate model. *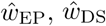* and *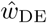* denote the total strength of connection between PV neurons and the soma of PCs, SOM neurons and the dendrites of PCs and the recurrence strength between PCs, respectively. The total connection strength is given by the product of the number of existing connections between two neuron types (or compartments) and the strength for individual connections. All parameters taken from Murayama et al. (2009). Note that we incorporated the gain factor present in Murayama et al. (2009) into the parameters to achieve unit consistency for all neuron types.

| parameter (unit) | value |
| --- | --- |
| $\Theta$ ($s^{-1}$) | 14 |
| $\lambda_E$ | 0.31 |
| $\lambda_D$ | 0.27 |
| $\hat{w}^{EP}$ | 0.7 |
| $\hat{w}^{DS}$ | 1.96 |
| $\hat{w}^{DE}$ | 0.42 |
| $c$ ($s^{-1}$) | 7 |
| $\Theta_c$ ($s^{-1}$) | 28 |

The firing rate dynamics of each interneuron is modeled by a rectified, linear differential equation (Wilson and Cowan, 1972):

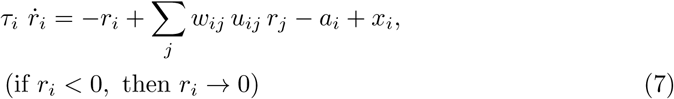

where *a*_*i*_ represents an adaptation variable, *w*_*ij*_ denotes the relevant synaptic weight onto the neuron, *u*_*ij*_ describes a synaptic facilitation variable and *x*_*i*_ denotes external inputs. The rate time constant *τ*_*i*_ was chosen to resemble the GABA_A_ time constant of approximately 10 ms for all interneuron types included. The weight matrix *W* (see Table 3) was chosen such that the relative connection strengths are consistent with experimental findings (Avermann et al., 2012; Pfeffer et al., 2013; Lee et al., 2013; Pi et al., 2013; Jiang et al., 2015). When we simulated mismatch neurons (see Figure 8),we set the SOM → PV connection to *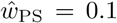* and tuned the connection from PV neurons to the somatic compartment of PCs in order to ensure a response only when the visual input is switched off and the (motor-related) prediction is switched on: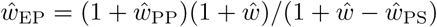. Note that all weights are scaled according to the number of existing connections, such that simulation results are robust to changes in the number of neurons in any of the cell classes.

**Table 3.** Connection strengths between neuron types. Entries in the same columns correspond to the same presynaptic neuron type, entries in the same row to the same postsynaptic neuron type. Given are the total connections strengths (absolute values, sign in simulations in line with neuron type – excitatory/inhibitory), which are the product of the number of existing connections between two neuron types (or compartments) and the strength for individual connections. The total recurrence strengths *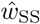* and *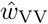* as well as the total mutual inhibition strengths *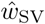* and *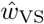* are varied in the simulations. Parentheses denote values that are only used when recurrence is introduced artificially.

|  | E | PV | SOM | VIP |
| --- | --- | --- | --- | --- |
| PV | 1 | 1.5 | 1.3 | - |
| SOM | 1 | - | $\hat{w}_{\text{SS}}$ | $\hat{w}_{\text{SV}}$ |
| VIP | 1 | - | $\hat{w}_{\text{VS}}$ | $\hat{w}_{\text{VV}}$ |

In contrast to PV neurons, both SOM and VIP cells show pronounced spike-frequency adaptation (Rudy et al., 2011; Gentet, 2012; Tremblay et al., 2016; Urban-Ciecko and Barth, 2016; Wamsley and Fishell, 2017), which is described by an adaptation variable *a*_*i*_,

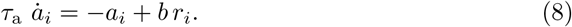

At constant neuronal activity *r*_*i*_, the adaptation variable *a*_*i*_ exponentially approaches the steady-state value *b r*_*i*_ with time constant *τ*_a_. For simplicity, if not otherwise stated, the adaptation strength *b* and time constant *τ*_a_ are the same for both cell types (*b ∈* [0, 2], *τ*_a_ = 100 ms). If adaptation is not present, we set the parameter *b* to zero.

Short-term facilitation is only modeled for SOM*→*VIP and VIP*→*SOM connections. The facilitation variable *u*_*ij*_ between neuron *j* and neuron *i* evolves according to the Tsodyks-Markram model (Tsodyks and Markram, 1997; Markram et al., 1998):

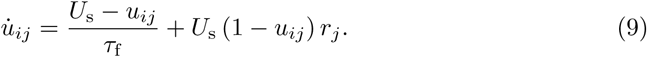

The facilitation variable *u*_*ij*_ ranges from 0 to 1 and represents the release probability, which changes according to the availability of calcium in the axon terminals. In the absence of presynaptic activity, the facilitation variable *u*_*ij*_ relaxes exponentially with time constant *τ*_f_ to a steady state *U*_*s*_, which represents the initial release probability. Presynaptic activity increases the facilitation variable *u*_*ij*_ by an amount proportional to *U*_s_. If not stated otherwise, the initial release probability *U*_s_ and facilitation time constant *τ*_f_ are equal for SOM and VIP neurons (*U*_s_ *∈* [0, 1], *τ*_f_ = 200 ms). When STF is not present, *u*_*ij*_ = 1 (or, equivalently, *U*_s_ = 1). In simulations where the strength of short-term facilitation is varied, we ensured comparability by scaling the weights *w*_*ij*_ by *U*_s_, thereby keeping constant the initial synaptic response after a long period of inactivity.

### External stimulation

To achieve physiologically reasonable activity levels, all neurons are stimulated with a time-independent background rate *x*_*i*_. PCs receive constant bottom-up input *x*_E_ at the soma and top-down feedback *x*_D_ at their dendrites. Additionally, VIP (in the full network setting) or SOM neurons (in the reference network) receive an external stimulus *x*_mod_ that was varied systematically to investigate the amplification properties of the microcircuit. If not indicated differently, all cells of the same neuron type are presented with an identical stimulus.

In the interneuron network, for the sake of comparability across different parameter settings, we always adjust the background inputs *x*_*i*_ such that the spontaneous activity (that is, at *x*_mod_ = 0) is equal to *r*_0_ = 3*/s* for all interneurons. In the non-WTA regime this can be achieved by

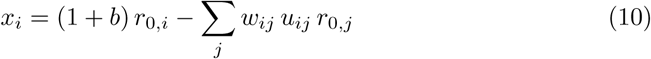

with *u*_*ij*_ = 1, if short-term facilitation is not present, or

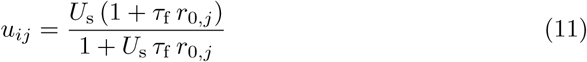

otherwise. In the full microcircuit comprising PC, PV, SOM and VIP cells, the external stimulation is set to *x*_PV_ = *x*_SOM_ = *x*_VIP_ = 3*/s* for the interneurons and *x*_E_ = 17.5*/s* and *x*_D_ = 21*/s* for the PC neurons, if not otherwise stated.

To characterize the dynamics of the system, we perform a frequency response analysis that measures the amplitude of the output signal as a function of the frequency 1*/T*. In that case, the weak external stimulus *x*_mod_ is expressed by a sine wave

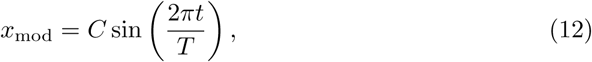

where *T ∈* [50, 600] ms. The amplitude of the output signal is normalized to the amplitude *C* of the input oscillation (*C* = 1s^-1^ in the simulations).

To investigate correlations, we stimulated SOM and VIP neurons with an input consisting of i) a constant component 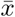 (calculated as before, see Equation 10), ii) individual noise and iii) noise that it shared among the neurons of the same type:

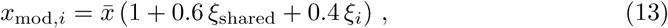

where the noise terms *ξ*_shared_ and *ξ*_*i*_ are drawn at each time *t* from Gaussian distributions with zero mean and unit variance. The shared component of the noise accounts for the strong correlations seen by Karnani et al. (2016). Furthermore, the number of cells per neuron type is increased in these simulations by a factor 5 to obtain reliable statistical estimates.

For the example firing rate traces (in Figure 5 and Figure S3 & S4), we stimulated SOM and VIP neurons with an input consisting of i) a constant component of 25*/s* and individual noise drawn at each time *t* from a Gaussian distribution with zero mean and SD of 5*/s*.

### Definition and mathematical derivation of the amplification index

To quantify the strength of amplification in our neural microcircuit, we introduce the amplification index,

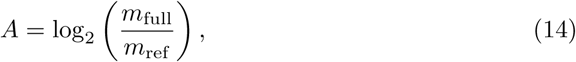

where *m*_full_ and *m*_ref_ denote the slope of the sigmoid function of the full and the reference network (see Figure 1 D or 2 B), respectively. These slopes represent the redistribution of somatic and dendritic inhibition upon a change in modulatory VIP input. Hence, the amplification index measures how much stronger the redistribution is when weak input passes through the SOM-VIP motif instead of directly through the SOM neurons.

The amplification index can be calculated analytically for the simplified linear network without PCs and short-term facilitation. To this end, we first derive the slope *m*_full_ from the mean-field population dynamics,

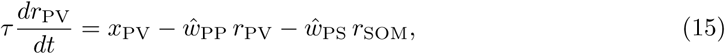

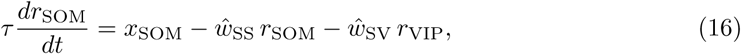

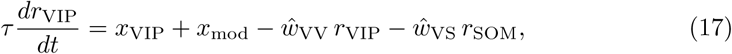

where 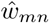 denotes the total weight from neuron type *n* onto *m*, and the rates *r*_*n*_ denote the mean-field population rate of neuron type *n*. For the sake of generality, we included the possibility of recurrent connections among both SOM and VIP neurons. Solving this set of equations for the steady-state population activity yields

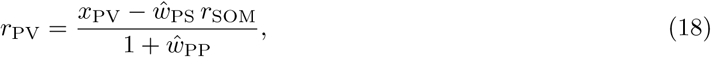

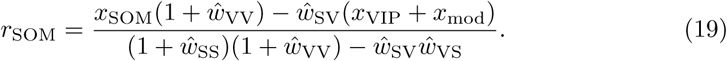

The slope *m*_full_ is given by the derivative of *r*_PV_ *-r*_SOM_ with respect to *x*_mod_,

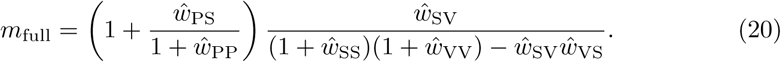

Similarly, the slope *m*_ref_ in the reference network can be derived from the mean field equations without VIP and PC neurons,

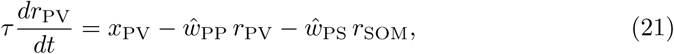

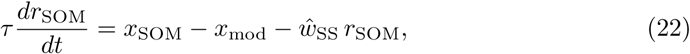

and is given by

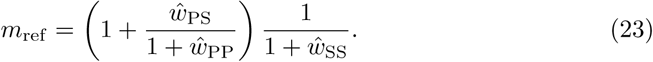

Finally, the amplification index is given by the ratio of the slopes,

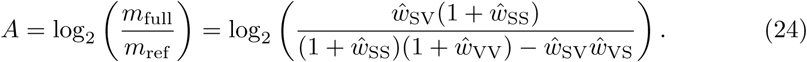

In the absence of recurrent inhibition within SOM and VIP populations, this expression simplifies to

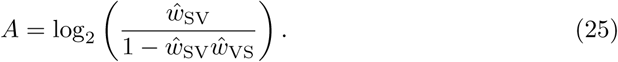

### Mathematical analysis of the computational repertoire of the SOM-VIP network

To analyze the computational repertoire of the SOM-VIP motif, we considered a network composed of SOM (hereafter only S) and VIP neurons (hereafter only V) that are mutually and fully connected. Again, the rectifying nonlinearity of the neurons is neglected.

### Computational states with recurrent inhibition within SOM and VIP neurons

In the absence of adaptation, the dynamics of the system is fully characterized by the change in rates. Each population comprises *n* cells, so the dynamics of the full network are described by a (2 *× n*)-dimensional system of linear differential equations,

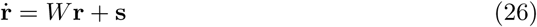

with 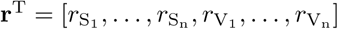 represents a vector of external stimuli and *W* is a block matrix that contains the connection weights.

By deriving the eigenvalue spectrum of the matrix *W* (see Appendix 1 for a detailed derivation), we can perform a bifurcation analysis. For the sake of simplicity, we suppose that the rate time constants, recurrence and mutual inhibition strengths are equal for both interneuron types: *τ*_S_ = *τ*_V_ = *τ, w*_SS_ = *w*_VV_ = *w*_r_ and *w*_SV_ = *w*_VS_ = *w*. By analyzing the sign and nature (complex or real) of the eigenvalues, we can define five dynamical regimes:

(i) All interneurons can be active and operate in an attenuation regime,

(ii) All interneurons can be active and operate in an amplification regime,

(iii) Winner-take-all (WTA) between SOM and VIP neuron population (strong competition, either all SOM or all VIP cells are silenced, neurons within the winning population are all active),

(iv) WTA in each population separately (exactly one VIP and one SOM cell remain active),

(v) Total WTA (only one single neuron in the whole network is active).

The transition between the attenuation and the amplification regime ((i) and (ii)) is determined by the condition that the amplification index, Equation 24, is equal to one (for the symmetric case 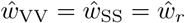 and 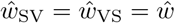). The transition to WTA regimes ((iv) and (v)) within each neuron population emerges when the total recurrence strength *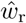* is greater or equal to the leak multiplied by the population size,

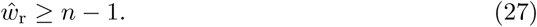

Moreover, the transition to a WTA regime between SOM and VIP neurons (regime (iii)) occurs when the total mutual inhibition strength *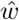* is larger than the sum of total recurrence 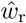 and leak,

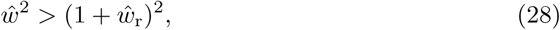

where 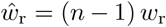 and 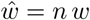. Finally, the pathological regime (v) of a total WTA requires that Equation 27 is fulfilled and that *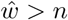*, i.e., condition 28 at the transition boundary to WTA within the two populations.

### Computational states with adaptation

In order to derive the qualitative changes in the bifurcation structure when adaptation (instead of recurrence) is present, we extended the rate-dynamics (see Equation 26) with linear differential equations describing the evolution of an adaptation current (cf. Equation 8). The (4*×n*)-dimensional state-vector **r** is now given by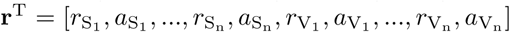 For the sake of simplicity and comparability, we suppose that the rate time constants, the mutual inhibition strength as well as the adaptation parameters are equal for SOM and VIP neurons: *τ*_S_ = *τ*_V_ = *τ, w*_SV_ = *w*_VS_ = *w, b*_S_ = *b*_V_ = *b* and *τ*_a,S_ = *τ*_a,V_ = *τ*_a_. This symmetry simplifies the derivation of the eigenvalues considerably (see Appendix 1).

The eigenvalue spectrum reveals that, in contrast to recurrence, adaptation does not lead to pathological states (see conditions (iv) and (v) in subsection above). Furthermore, depending on the sign and nature of the eigenvalues, we find four dynamical regimes:

(i) All interneurons can be active and operate in an attenuation regime (Figure 5A, region a),

(ii) All interneurons can be active and operate in an amplification regime (Figure 5A, region b),

(iii) Switch regime (Figure 5A, region c): WTA between SOM and VIP neuron population when total mutual inhibition strength is larger than the sum of the adaptation strength and the leak:

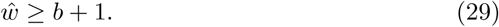

(iv) Oscillatory switch regime (Figure 5A, region d), in which SOM and VIP cells alternate between active and inactive states. It requires two conditions: First, adaptation must be stronger than the difference of total mutual inhibition and leak. Second, this total reciprocal inhibition must be larger than leak and ratio of rate and adaptation time constant:

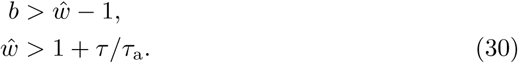

The derivation of the amplification index *A* for a network with adaptation is analogous to the derivation with recurrent inhibition, and merely requires to replace the total recurrence strength 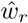 by the strength *b* of adaptation. Hence, the transition between the attenuation and the amplification regime ((i) and (ii)) is determined by this minor modification of Equation 24.

In the oscillatory switching regime, some of the eigenvalues of the dynamical system are complex. An approximation of the oscillation frequency can then be derived from their imaginary part:

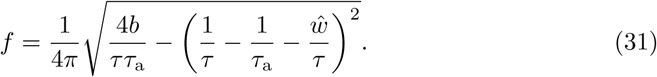

## Simulation details

All simulations were performed in customized Python code written by LH. Differential equations were numerically integrated using a 2^nd^-order Runge-Kutta method with a maximum time step of 0.05 ms. Neurons were initialized with *r*_*i*_(0) = 0 Hz, *a*_*i*_(0) = 0 Hz (if adaptation was modeled) and *u*_*ij*_(0) = *U*_s_ (if STF was present) for all *i*. Source code will be made publicly available upon publication.

## Acknowledgments

We are grateful to Laura Bella Naumann and Filip Vercruysse for a critical reading of the manuscript. The project is funded by the German Federal Ministry for Education and Research, FKZ 01GQ1201.

## Appendix 1

### A. Mathematical analysis of the SOM-VIP motif

In order to characterize the computational repertoire of the SOM-VIP motif, we performed an extensive mathematical analysis of a simplified model that comprises only SOM and VIP cell populations. Moreover, we neglected the rectifying nonlinearity that ensures that firing rates remain positive. We will start with networks that take into account either recurrent connections among both SOM and VIP neurons or adaptation, but ignore short-term plasticity. In these cases, the circuits are described by linear dynamical systems and therefore allow detailed bifurcation analyses. The derivations provide analytical conditions for the parameter boundaries between different computational states of the SOM-VIP motif. We then continue to derive approximations for these boundaries for the nonlinear case that includes short-term plasticity.

#### A.1 Bifurcation analysis: Recurrent inhibition within the SOM/VIP populations

We consider a network composed of SOM (hereafter only S) and VIP neurons (hereafter only V) that are mutually and fully connected. Furthermore, we assume that each population comprises *n* cells. The circuit dynamics can then be described by

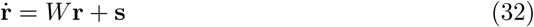

with 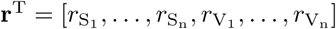. **s** represents a vector of external stimuli and *W* is a block matrix of the form

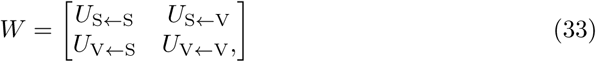

which contains the connection weights. When recurrence is included, the (*n × n*) blocks have the following

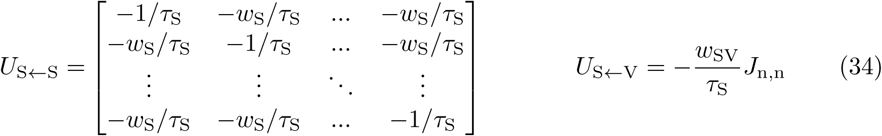

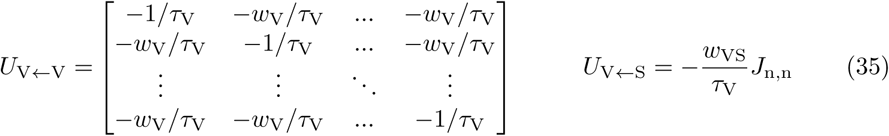

where *J*_n,n_ is a squared matrix of dimension *n* with all entries equal one. Note that autapses were not included and that we chose the convention that the weight parameters *w*_*i*_ are positive.

The dynamical properties of the circuit are described by its eigenvalues, which are given by the zero crossings of the characteristic polynomial of the matrix *W*,

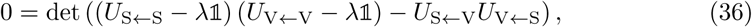

where 1 denotes the *n*-dimensional identity matrix. Resolving the determinant yields the following condition for the eigenvalues:

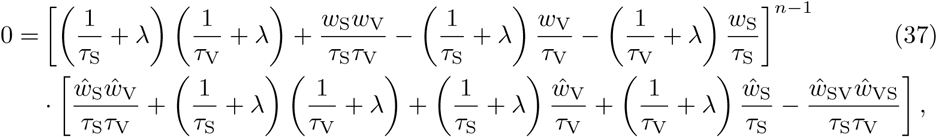

with the total recurrence strengths 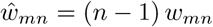 and the total mutual inhibition strengths 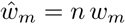 Consequently, the eigenvalues of the linear system are

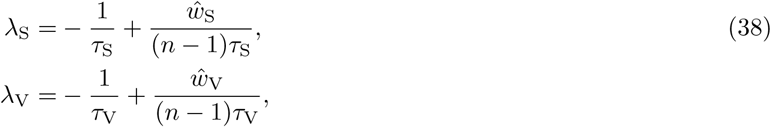

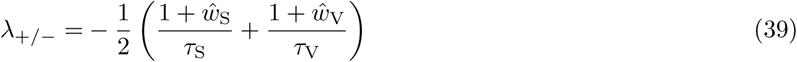

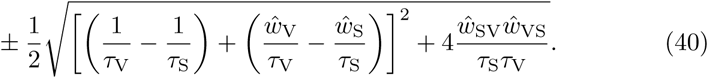

While the first and second eigenvalues *λ*_*S*_, *λ*_*V*_ have an algebraic and geometric multiplicity of (*n -* 1) and describe the dynamics of rate inhomogeneities within the two populations, the eigenvalues *λ*_+_ and *λ*_*-*_ have an algebraic and geometric multiplicity of one and determine the dynamics of the population rates of the two interneuron types. Because 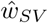 and 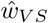 are positive, all eigenvalues are real, suggesting the absence of oscillations. For the sake of simplicity, we now analyze a symmetric situation, in which *τ*_S_ = *τ*_V_ = *τ, 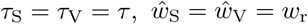* and *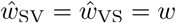* By analyzing the sign and the nature (complex or real) of the eigenvalues, we can define five dynamical regimes:

i. All interneurons can be active and operate in an attenuation regime (no competition, all eigenvalues negative, weak mutual inhibition),
ii. All interneurons can be active and operate in an amplification regime (no competition, all eigenvalues negative, strong mutual inhibition),
iii. WTA between SOM and VIP neuron population (either SOM or VIP cells win; *λ*_+*/-*_ *>* 0, *λ*_*S/V*_ *<* 0),
iv. WTA in each population separately (one VIP and one SOM cell survive, *λ*_+*/-*_ *<* 0, *λ*_*S/V*_ *>* 0 positive),
v. Total WTA (only one single neuron is active, all eigenvalues positive).

The transition between the attenuation and the amplification regime ((i) and (ii)) is determined by the condition that the amplification index Equation 24 is equal to one (for the symmetric case 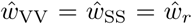 and *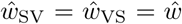* The transition to a WTA regime within each neuron population emerges when *λ*_S_ = *λ*_V_ = *λ* ≥0, yielding the condition

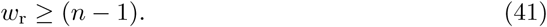

Moreover, the transition to a WTA regime between SOM and VIP neurons occurs when *λ*_+_ *≥* 0, which yields

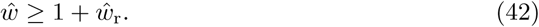

#### A.2 Bifurcation analysis: Adaptation

We extend the rate dynamics (see Equation 32) with linear differential equations describing the evolution of an adaptation current in order to derive qualitative changes in the bifurcation structure when adaptation (instead of recurrent inhibition) is present. The (4*×n*)-dimensional state vector **r** now includes the adaptation variables and is given by 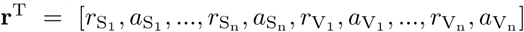. Again, the dynamical system can be written in the format of Equations 32 and 33, but the matrices *U*_m*?*n_ (*m, ∈n* {*S, V* }) now themselves become block matrices:

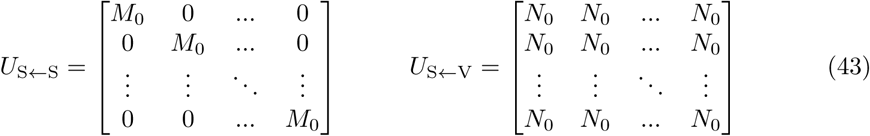

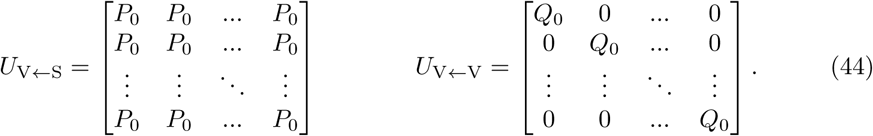

Here, *M*_0_, *N*_0_, *P*_0_ and *Q*_0_ are (2 *×* 2) matrices given by

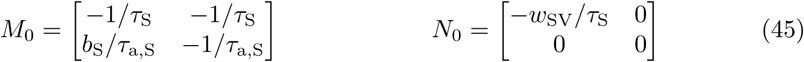

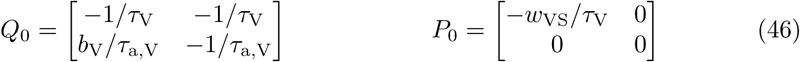

with the adaptation strengths *b*_S_ and *b*_V_, as well as adaptation time constants *τ*_a,S_ and *τ*_a,V_. As the block matrices do not commute, the eigenvalue condition for the characteristic polynomial is now given by (Silvester, 2000)

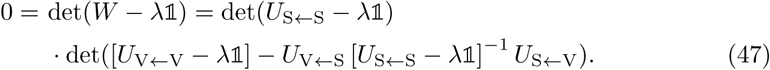

After some straightforward linear algebra, the eigenvalues are given by the zeros of the following equation:

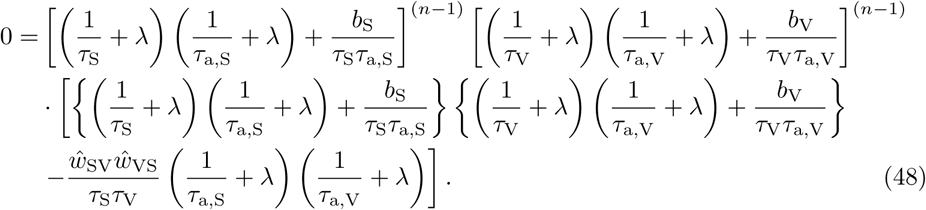

For the sake of simplicity and comparability, we again consider the symmetric case *τ*_S_ = *τ*_V_ = *τ, w*_SV_ = *w*_VS_ = *w, b*_S_ = *b*_V_ = *b* and *τ*_a,S_ = *τ*_a,V_ = *τ*_a_. This simplifies the derivation of the eigenvalues considerably. We obtain three pairs of eigenvalues

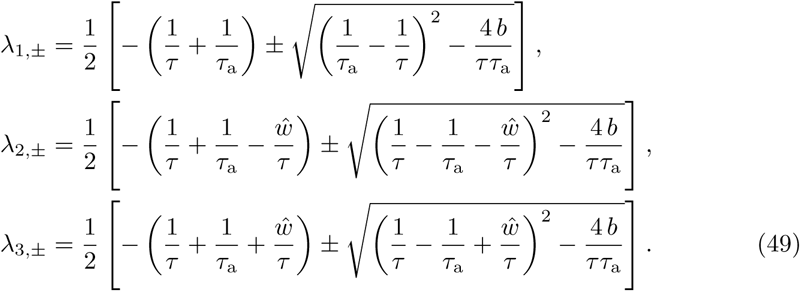

The first two eigenvalues (*λ*_1,+_ and *λ*_1,*-*_) have both algebraic and geometric multiplicity of (2*n-*2) and define the rate and adaptation dynamics of rate inhomogeneities within the two interneuron populations. Both eigenvalues are strictly negative for all parameters possible. Hence, in contrast to recurrence, adaptation does not lead to pathological states in which a single cell within one population silences the others (see conditions and (v) in section above). The last four eigenvalues have algebraic and geometric multiplicity of one and describe the interaction between the two populations. It can be easily seen that Re(*λ*_3,*±*_) *<* 0, so that instabilities – like WTA regimes – can only arise from the two eigenvalues *λ*_2,*±*_. Depending on the sign and nature of these eigenvalues, we find four dynamical regimes:

(i) All interneurons can be active and operate in an attenuation regime (all eigenvalues real and negative, weak mutual inhibition),

(ii) All interneurons can be active and operate in an amplification regime (all eigenvalues real and negative, strong mutual inhibition),

(iii) WTA between SOM and VIP neuron population, described by the condition

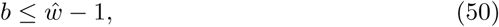

(iv) An oscillatory WTA regime (osc WTA) in which SOM and VIP neurons alternate between active and inactive states. It depends on two conditions

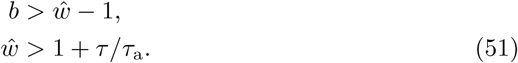

The derivation of the amplification index *A* for a network with adaptation is analogous to the derivation with recurrent inhibition, and merely requires to replace the total recurrence strength 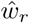 by the strength *b* of adaptation. Hence, the transition between the attenuation and the amplification regime ((i) and (ii)) is determined by this minor modification of Equation 24.

Strictly speaking, the last regime (iv) has to be separated into two subregimes, in which the two eigenvalues *λ*_2,*±*_ either (a) form a complex conjugate pair with positive real part or (b) are both real and positive. In the former regime, an approximation of the oscillation frequency *f* can be obtained from the imaginary part of the eigenvalues:

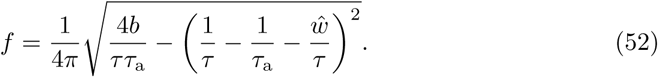

The oscillation frequency follows a square root function in *b* with a scaling factor determined by *τ*_a_, and an offset controlled by both 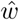 and *τ*_a_. The parameter regime (a), in which an approximation for the oscillation frequency can be obtained from a linear analysis is limited by the condition

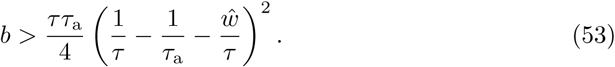

In the second regime (b), the circuit nevertheless displays oscillations when simulated. These rely on the rectifying nonlinearity for the firing rates, so their frequency cannot be calculated within the present linear analysis.

#### A.3 Bifurcation analysis: Adaptation and short-term facilitation

When synaptic adaptation (STF) is included in the model, the dynamical system equations become nonlinear and mathematically intractable. Thus, we can only derive approximate expressions for the transitions between the computational regimes (see (i) to (iv) in section above).

In zero-order approximation, we assume that the facilitation variable *u*_*ij*_ can be replaced by its steady-state value *u*_*∞*_, for which we derive an expression below. The boundaries given by Equations 50 and 51 can then be replaced by

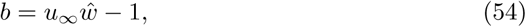

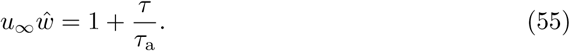

These approximations allow to deduce the qualitative changes in the transition boundaries. As *u*_*∞*_ *≥* 1, the slope of Equation 54 increases, leading to an enlarged WTA regime (i.e., switch-like state). Moreover, the transition to an oscillatory switch is pushed towards smaller mutual inhibition strengths and requires stronger adaptation (cf. Equations 54- 55).

To qualify the regime boundaries, we have to derive an expression for the steady-state value *u*_*∞*_ as a function of the parameters *b* and 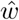. To this end, we identify a self-consistent solution of the equations for *u*_*∞*_ and the steady state of the population rates of the two interneuron types.

The steady state of the population rates 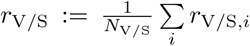 is given by the conditions

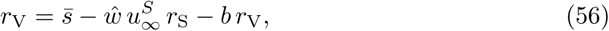

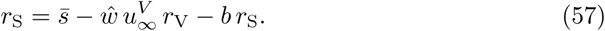

Here, 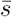 denotes the averaged input. 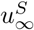 and *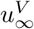* represent the steady-state facilitation variable for SOM*→*VIP and VIP*→* SOM, respectively, which are given by

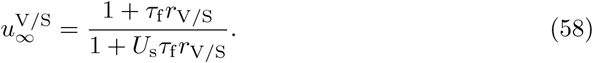

To express *u*_*∞*_ in terms of *b* and *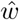*, we have to resolve Equation 57 for the population rates and insert these expressions into Equation 58. Because the resulting equations are lengthy and non-informative, we consider two approximations. In the non-WTA regime, as intrinsic and synaptic parameters as well as the external stimulation are taken to be equal, we assume that the population firing rates of SOM and VIP neurons are approximately equal on average *r*_V_ *≈ r*_S_ *≡r*. This leads to the following expression for the population rates *r* in terms of the system parameters:

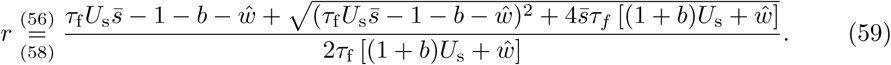

By inserting Equations 58 and 59 into Equation 55, we obtained a good approximation of the transition boundary between the amplification regimes and the oscillatory switch state.

For the transition to a WTA regime, the assumption of equal rates is violated. To derive an expression for this transition boundary, we therefore replace the assumption of equal rates by an assumption of a WTA state, in which one of the rates is zero. Because all parameters are assumed to be symmetric, we can assume without loss of generality that the VIP population is the winner. We can then obtain conditions for the boundary by setting *r*_*S*_ = 0 in Equations 56 and 57 and inserting Equation 58 into Equation 57:

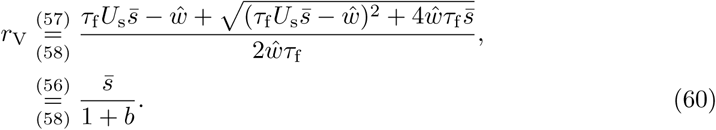

Solving this equation for the adaptation strength *b*, we obtain

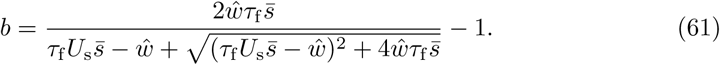

This approximation predicts the transition to the WTA regime qualitatively and quantitatively very well.

For deriving the transition boundary between attenuation and amplification, we note that the amplification threshold *A* = 1 is satisfied when the slopes for the full network and the reference network are equal,

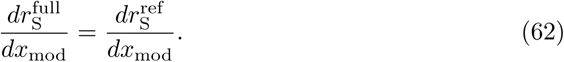

The steady state equation for the population rates of the SOM neurons in the reference and full network, respectively, are given by

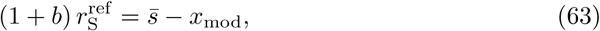

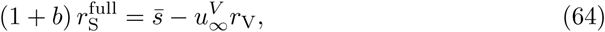

where 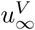 is given in Equation 58. The population rate *r*_V_ of the VIP neurons in the full network obeys

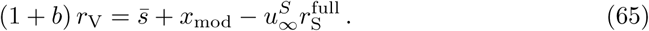

Taking the derivative of the steady-state Equations 63 and 64 for full and reference network with respect to *x*_mod_ yields

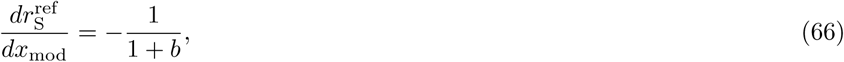

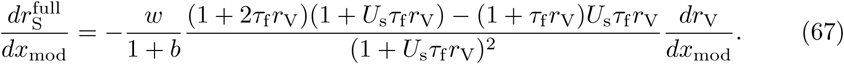

By taking the derivative of Equation 65, we can express 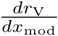 as a function of 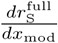 and 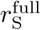. Furthermore, we assume again that *r*_V_ ≈ *r*_S_ ≡ *r* (cf. Equation 59). Solving for 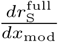, the condition 62 finally yields

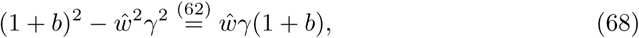

With

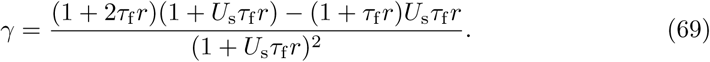

The combination of Equations 68, 69 and 59 provide an accurate prediction of the amplification threshold when STF and adaptation are present.

## B Supporting Figures

**Figure S1.**
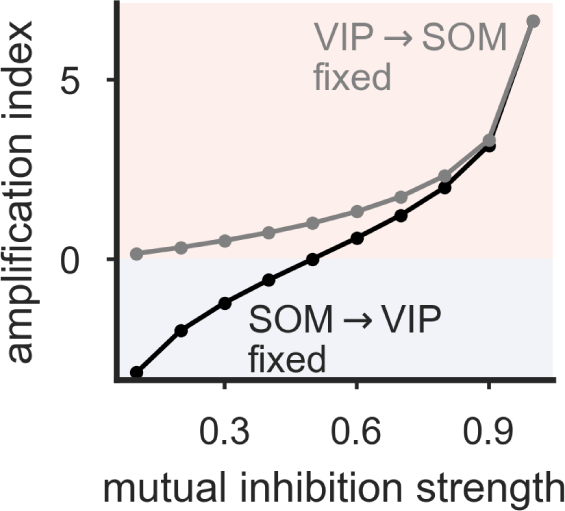
Asymmetric mutual inhibition strengths for SOM and VIP neurons also enhances the amplification index. When one of the connections, VIP*→*SOM 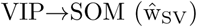 or SOM*→*VIP 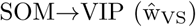, is kept constant, increasing the respective other weight leads to a strengthening of the amplification. Fixed weight was set to *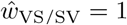*.

**Figure S2.**
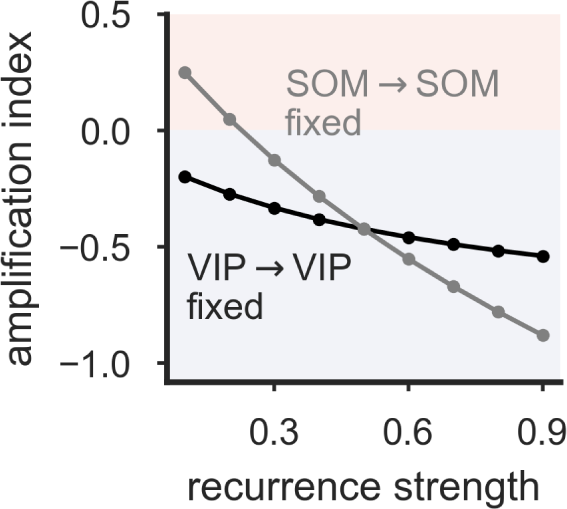
Asymmetric recurrence strengths for SOM and VIP neurons also reduce the amplification index. When one of the recurrent connections, VIP*→*VIP 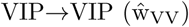 or SOM*→*SOM 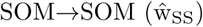, is kept constant, increasing the respective other weight leads to a decrease of the amplification index. Fixed weight was set to 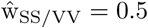 Mutual inhibition strength *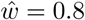*.

**Figure S3.**
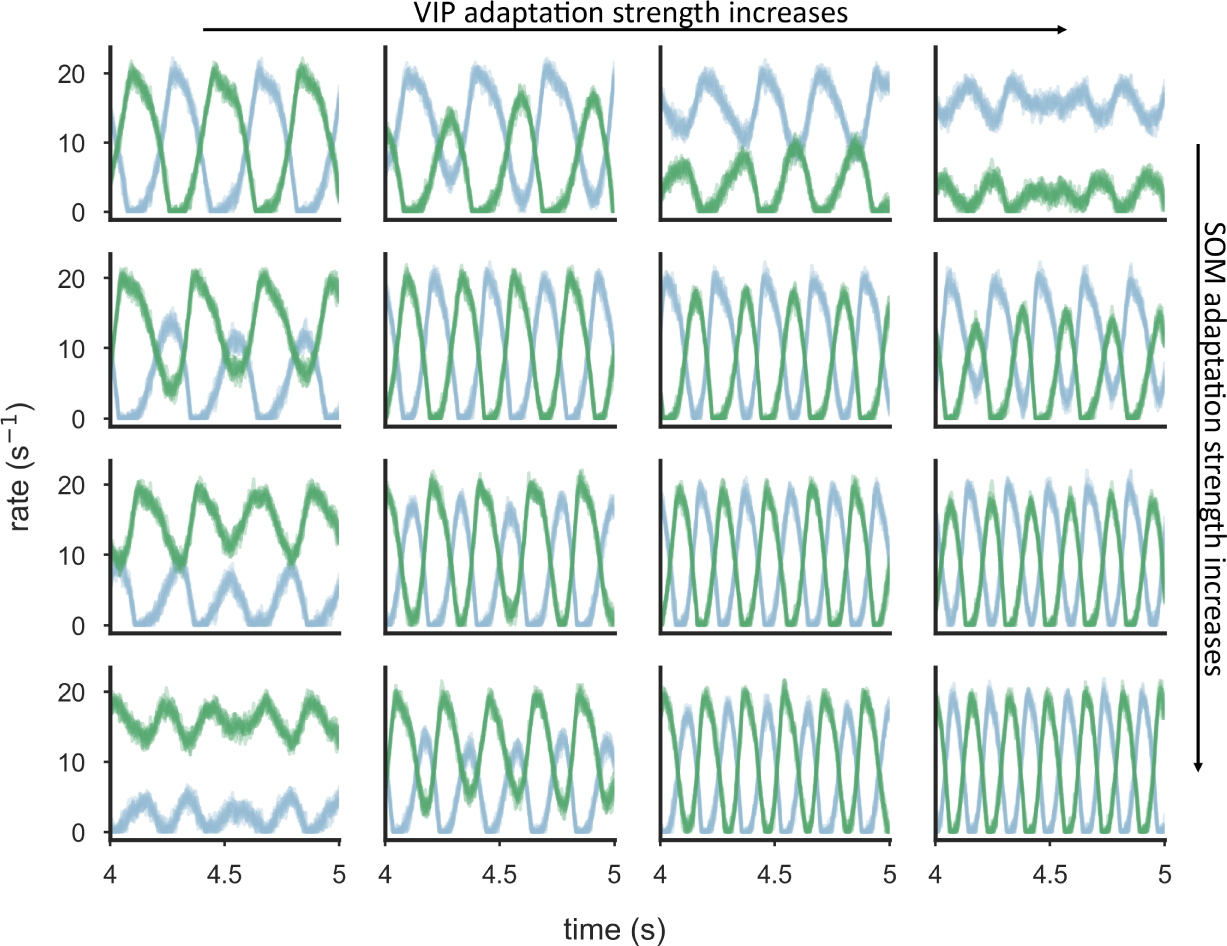
Oscillations also arise for asymmetric adaptation strengths in SOM and VIP neurons, with altered firing rate and oscillation frequency. Firing rate traces for SOM (blue) and VIP (green) neurons for a range of adaptation strengths (*b*_S*/*V_ *∈ {*0.4, 0.6, 0.8, 1*}*). Off-diagonal plots correspond to asymmetric adaptation strengths. Mutual inhibition strength *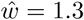*, adaptation time constants *τ*_*a*_ = 50 ms.

**Figure S4.**
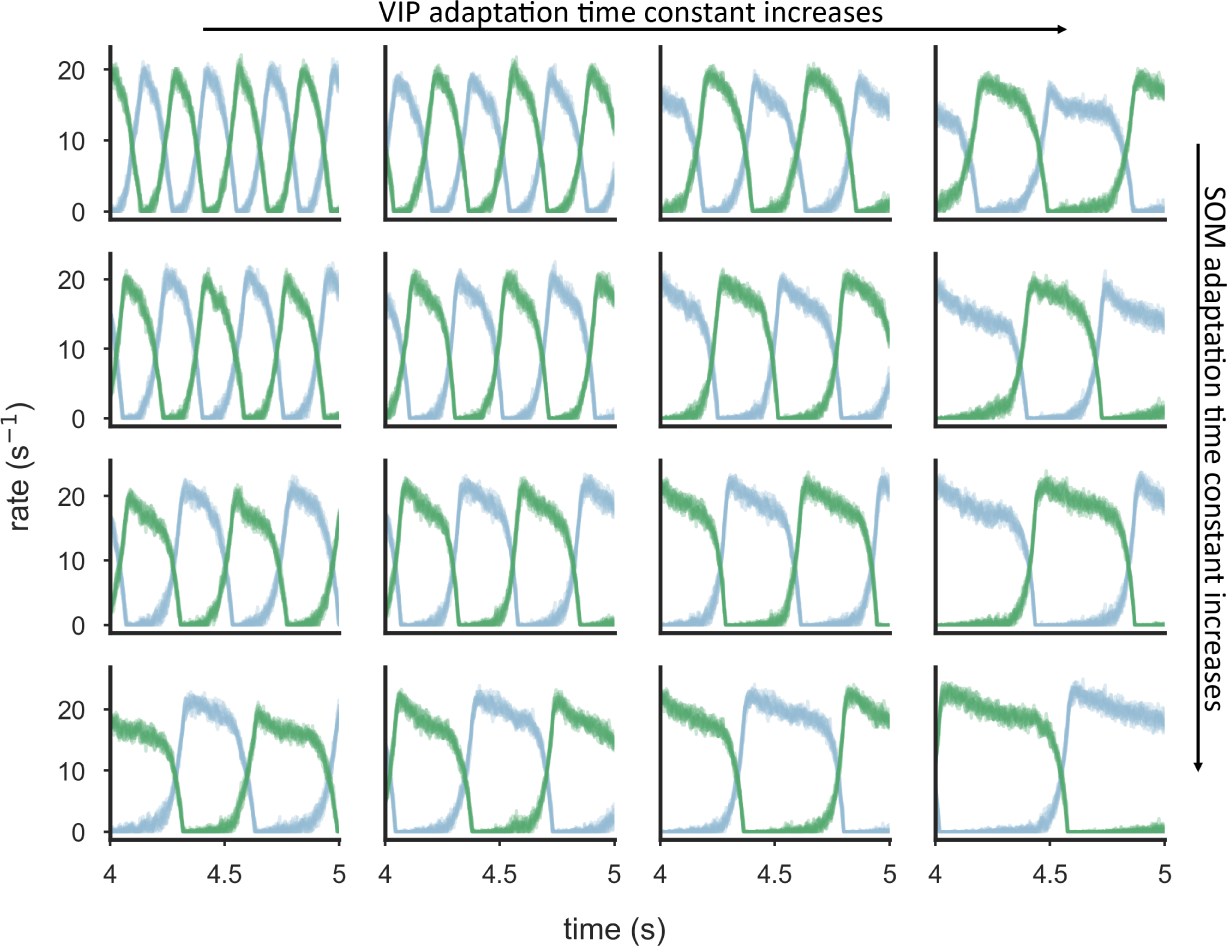
Asymmetric adaptation time constants for SOM and VIP neurons lead to different duration of active and inactive periods. Firing rate traces for SOM (blue) and VIP (green) neurons for a range of adaptation time constants (*τ*_*a,S/V*_ *∈* {50, 100, 200, 400} ms). Larger adaptation time constants cause longer active states. Mutual inhibition strength *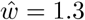*, Adaptation strength *b* = 0.5.

**Figure S5.**
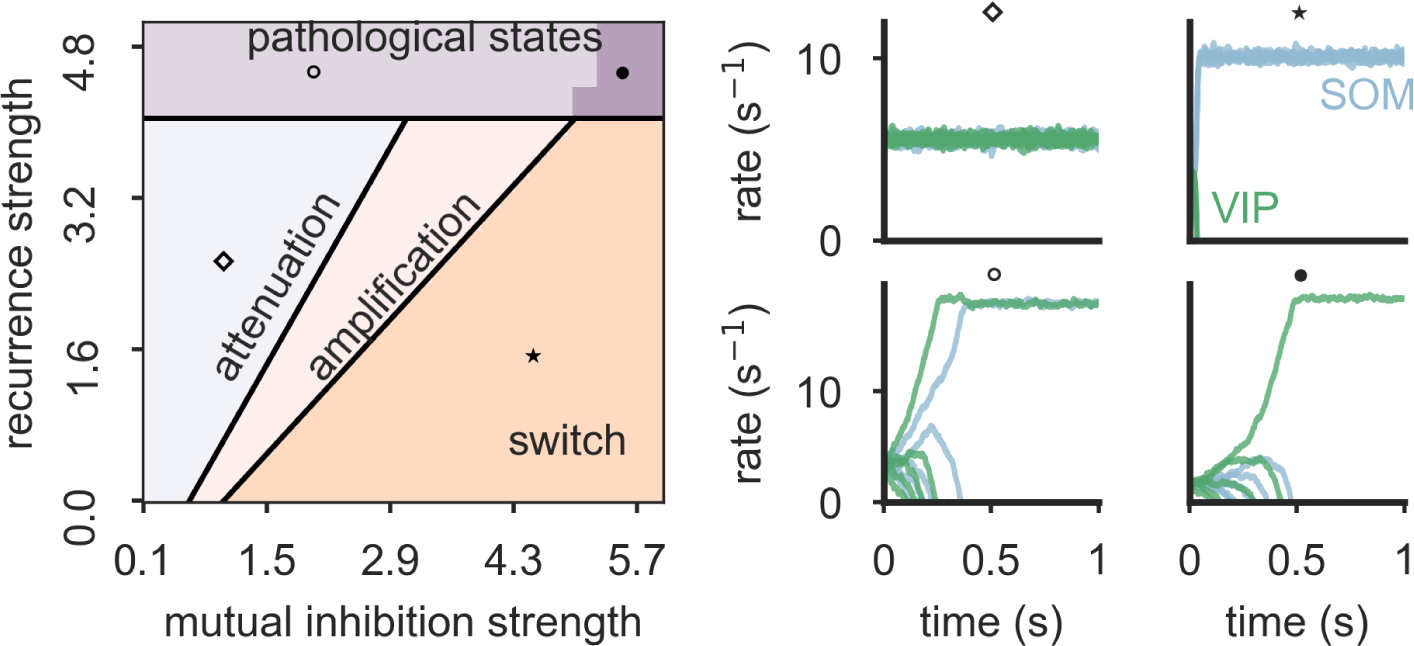
Dynamical states of the SOM-VIP motif with recurrence. Left: Bifurcation diagram reveals distinct operation modes: all interneurons are active (divided into amplification and attenuation regime), winner-take-all (WTA) regime leading to a switch, and two pathological states (WTA in each population separately and total WTA). Regime boundaries (black lines) are obtained from a mathematical analysis (see Appendix). Right: Example firing rate traces for all SOM (blue) and VIP (green) neurons for four network settings (see markers) taken from the bifurcation diagram. *N*_SOM_ = *N*_VIP_ = 5.

**Figure S6.**
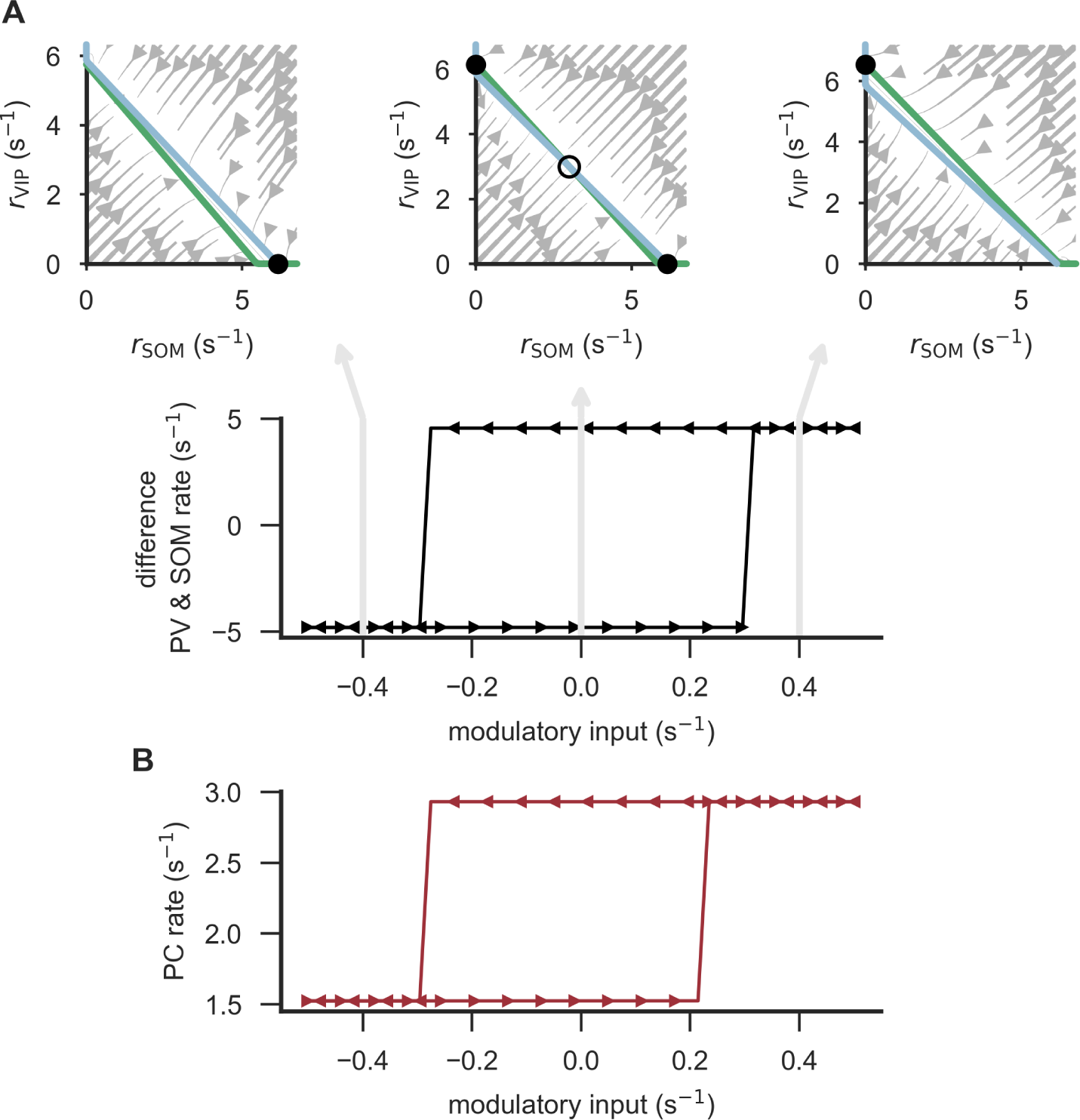
WTA between SOM and VIP neurons causes hysteresis in the reduced interneuron network and full microcircuit. **(A)** Example phase planes (top) for three distinct values of the modulatory input (cf. bottom). The intersection points of SOM-(blue) and VIP-nullcline (green) correspond to the fixed points that are either stable (filled circle) or unstable (open circle). The vector field shows the direction and strength of flow. In a WTA regime, the network exhibits bistability for a range of modulatory input values, leading to hysteresis (bottom). Mutual inhibition strength *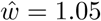* **(B)** Same as above for the full microcircuit. PC rate exhibits two stable states for a range of modulatory input. The steady-state activity depends on the initial state. Mutual inhibition strength *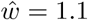*.

